# Transcriptome analysis of the winter tick (*Dermacentor albipictus*) reveals sex-specific expression patterns

**DOI:** 10.1101/2025.02.17.638687

**Authors:** Rhodri T.M. Edwards, Igor Antoshechkin, Eddie Hill, Michael W. Perry, Pia U. Olafson, Perot Saelao, Kimberly H. Lohmeyer, Omar S. Akbari

**Affiliations:** School of Biological Sciences, Department of Cell and Developmental Biology, University of California, San Diego, La Jolla, CA 92093; Division of Biology and Biological Engineering (BBE), California Institute of Technology, Pasadena, CA91125, USA; Livestock Arthropod Pests Research Unit, USDA-ARS, Kerrville, TX 78028, USA; Veterinary Pest Genetics Research Unit, USDA-ARS, Kerrville, TX 78028, USA

**Author notes:** To whom correspondence should be addressed: Omar S. Akbari, Ph.D., School of Biological Sciences, Department of Cell and Developmental Biology, University of California, San Diego, La Jolla, CA 92093, USA.

**Keywords:** RNAseq, transcriptome, Winter Tick, *D.albipictus*, vector control, pgSIT, *doublesex*

## Abstract

The winter tick, *Dermacentor albipictus*, is a significant ectoparasite of ruminants across North America, posing health risks to wildlife and occasionally humans. Despite its ecological importance, limited genomic resources exist for this species. This study provides the first comprehensive transcriptome analysis of *D. albipictus*, focusing on early-stage embryos, sexed adults, dissected ovaries, and dissected male reproductive systems. Differential gene expression analyses revealed significant sex-biased expression patterns, and functional annotations identified candidate genes involved in sex determination. Notably, we identified the first documented case of sex-specific splicing of a doublesex-like gene in chelicerates, a mechanism previously thought to be absent in this group. This discovery suggests that ticks may share more insect-like features of sexual differentiation, with implications for understanding the evolution of sex determination pathways in arthropods These transcriptome data serve as a critical resource for understanding the biology of *D. albipictus* and will facilitate the development of novel genetic control strategies.

## Introduction

The winter tick, *Dermacentor albipictus* (Family: Ixodidae), is a species of hard tick that is primarily distributed across North America (Chenery et al. 2023). It is an ectoparasite that primarily infests ruminants, including moose (Samuel 1989; Calvente et al. 2020a), elk (Calvente et al. 2020b), white-tailed deer (Calvente et al. 2020c; Machtinger et al. 2021), domestic bovids and equines (Sundstrom et al. 2021), and domestic cats and dogs (Duncan et al. 2020a). Winter tick infestations can cause skin conditions like alopecia and dermatitis (Duncan et al. 2020b; McLaughlin and Addison 1986). They may also be able to serve as vectors for atypical bacterial pathogens: studies have detected *Anaplasma spp.* (Ewing et al. 1997; Baldridge et al. 2009a), Francisella-like endosymbionts (Baldridge et al. 2009b), and *Borrelia burgdorferi* (Magnarelli et al. 1986; Kocan et al. 1992) in winter ticks.

While winter ticks primarily parasitize non-human mammals, they are not without risk to humans. A review of hard tick bites in humans showed that there have been 465 (0.2%) bites from *D. albipictus* in the United States compared to 158,008 (67.3%) from *Ixodes scapulari*s, which is a major vector for multiple human infections (Eisen 2022). Winter ticks largely affect hunters exposed to moose and deer (Rand et al. 2007). In humans, *D. albipictus* has been implicated in the transmission of *Babesia duncani*, a potentially fatal pathogen that can cause Babesiosis (Swei et al. 2019a).

Tick-borne zoonotic diseases are second only to mosquito-borne diseases in incidence and variety of pathogens (de la Fuente et al. 2008). Alarmingly, tick-host interactions are expected to increase in number in response to global climate change (Swei et al. 2019b; Pouchet et al. 2024; Gilbert 2021). Current control strategies primarily focus on personal protection, landscape and vegetation management, and acaricide use (Eisen 2021). These are not without risk, however, as the indiscriminate use of acaricides is partly responsible for the increase in reports of multi-resistant tick strains (Agwunobi, Yu, and Liu 2021). There is a need to identify new methods for control, including new acaricides, vaccine targets, and genetic tools.

Currently, 16 complete reference genomes are available for Ixodid tick species via NCBI (Jia et al. 2020; Guerrero et al. 2021; Chou et al. 2023; De et al. 2023), including a chromosome-level assembly of the *D. albipictus* genome (GCA_038994185.2). Tissue-specific transcriptome studies in ticks have analyzed expression in dissected tissues that have significance to feeding and reproduction: midgut (Xu et al. 2016; Omondi et al. 2023), salivary glands (Pienaar et al. 2021; Giachetto et al. 2019; Schwarz et al. 2013), male reproductive system (Sonenshine et al. 2011a), and ovaries (Cossío-Bayúgar et al. 2024; Zhao, Qu, and Jiao 2021). However, time points and tissues that would be most useful for genetic tool development are lacking.

In this study, we analyzed the transcriptome of *D. albipictus* in early-stage embryos (when constructs are typically introduced for transgenesis). We also analyzed the transcriptome of sexed adults, dissected ovaries, and dissected male reproductive systems in an effort to identify sex-specific expression and splicing of genes. We observed greater numbers of genes expressed in male-linked samples compared to female-linked samples, identified the most abundant gene ontologies, and explored a putative *doublesex* gene as a potential target for genetic control. The results advance our understanding of tick biology, development, and sex determination and also serve as a useful resource for the development of novel genetic control strategies that use sex-based sorting, such as precision-guided sterile insect technique (pgSIT).

## Results

### Overview of Sequencing, Gene Expression, and Principal Component Analysis

Transcriptome sequencing was performed to provide an overview of gene expression in *D. albipictus* unfed males (ufMale), unfed females (ufFem), the male reproductive system (Mrs), ovaries preoviposition (Ov-preov), and 0-6hr old embryos (embr_0-6h). Unfed adult samples were collected to provide comparisons between males and the pre-vitellogenic phase females (i.e. before oogenesis has occurred). Mrs and Ov-preov samples were collected to investigate differences in germline-specific gene expression in males and females. We generated an extensive dataset comprising 1.2 billion paired end 150 base reads with an average of 79.95 million reads per sample and a total yield of 332.8 Gb. 92.85% of reads were mapped to the *D. albipictus* genome with 85.95% being uniquely mapped (**Table 1**; **Table 2**).

**Table 1.**
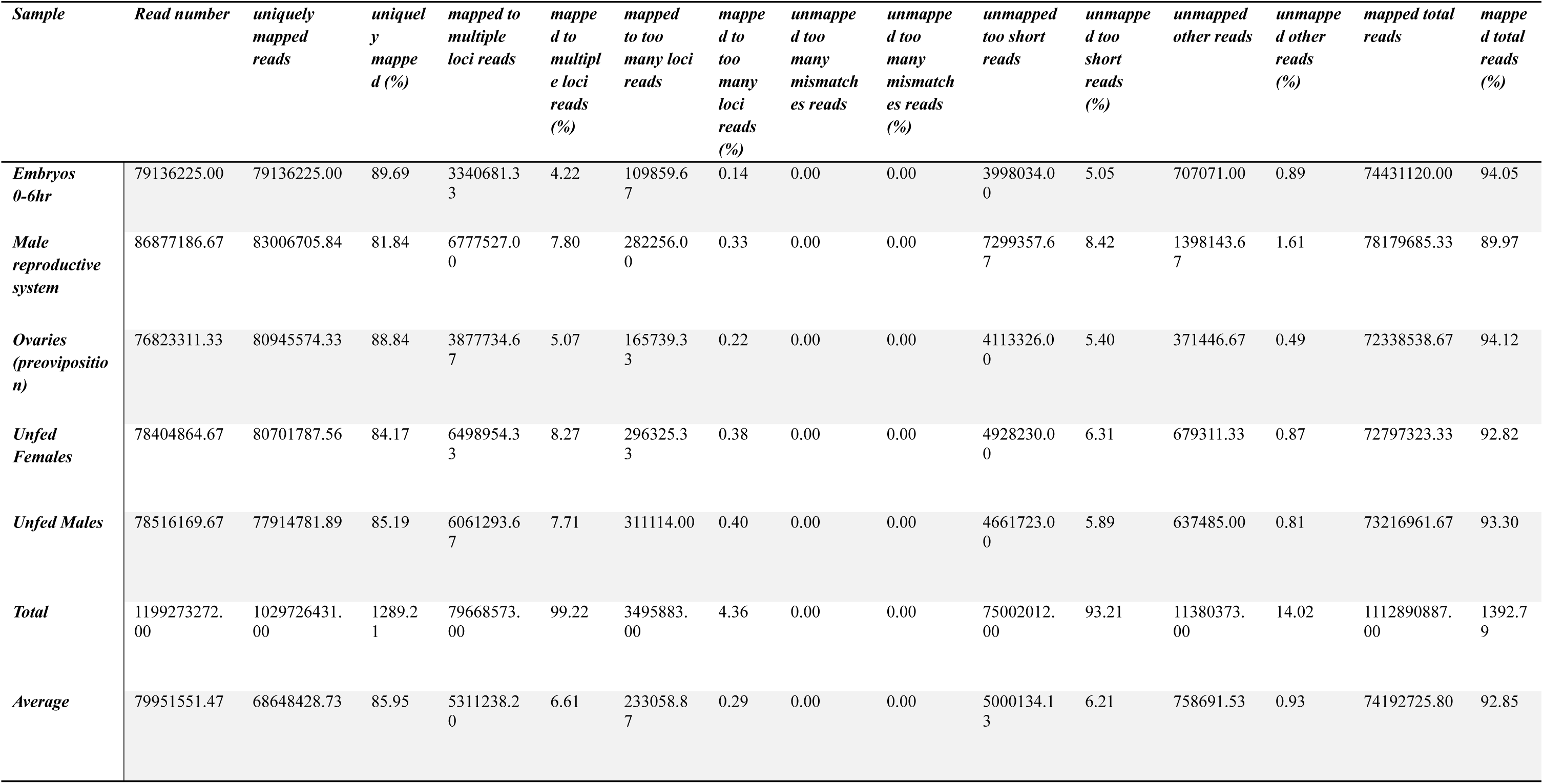
Summary of Illumina sequencing reads collected for *D.albipictus* samples (average of biological replicates), showing total reads, mapped and unmapped reads.

**Table 2.** Illumina sequencing mapping statistics for *D.albipictus* samples, showing total reads, mapped and unmapped reads, and read length. [mapping_stats_finalv2.xlsx](https://docs.google.com/spreadsheets/d/1E2Q7VB83m1FDYCRgqk0l9HxtrrvhA7Eq/edit?gid=358397184#gid=358397184)

Because no annotations were available for the GCA_038994185.2 genome, transcripts were assembled with StringTie for each of the sequenced samples individually and merged to generate a unified set of gene models comprising 22,680 genes and 47,811 transcripts. To provide a general overview of gene expression levels, the mean number of detected genes at FPKM > 1 for each sample type was determined and ranged from 11,336 in ufFem to 15,834 in ufMale. Embr_0-6h, ov-preov and Mrs samples expressed intermediate numbers of genes of 11,763, 11,882, and 14,986, respectively (**Figure 1**; **Table 1**; **Table 3**). Hierarchical clustering and principal component analysis on TPM values were used to visualize relationships between samples and showed tight grouping of replicates with well-separated sample types, as expected. (**Figure 2**).

**Figure 1.**
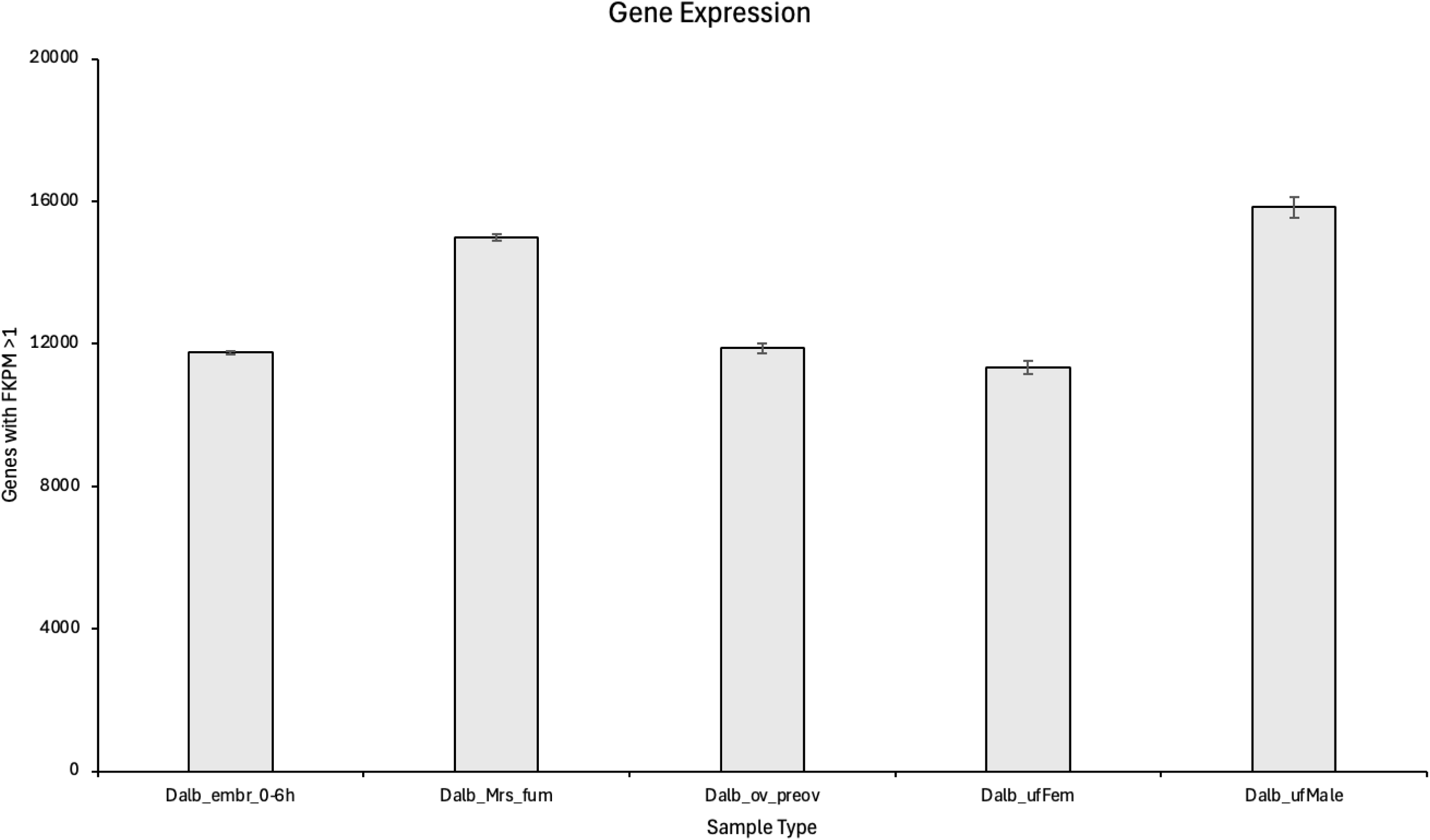
Number of expressed genes (FKPM >1) for all *D.albipictus* sample types. Error bars show standard deviation around the mean for biological replicates of each sample.

**Figure 2.**
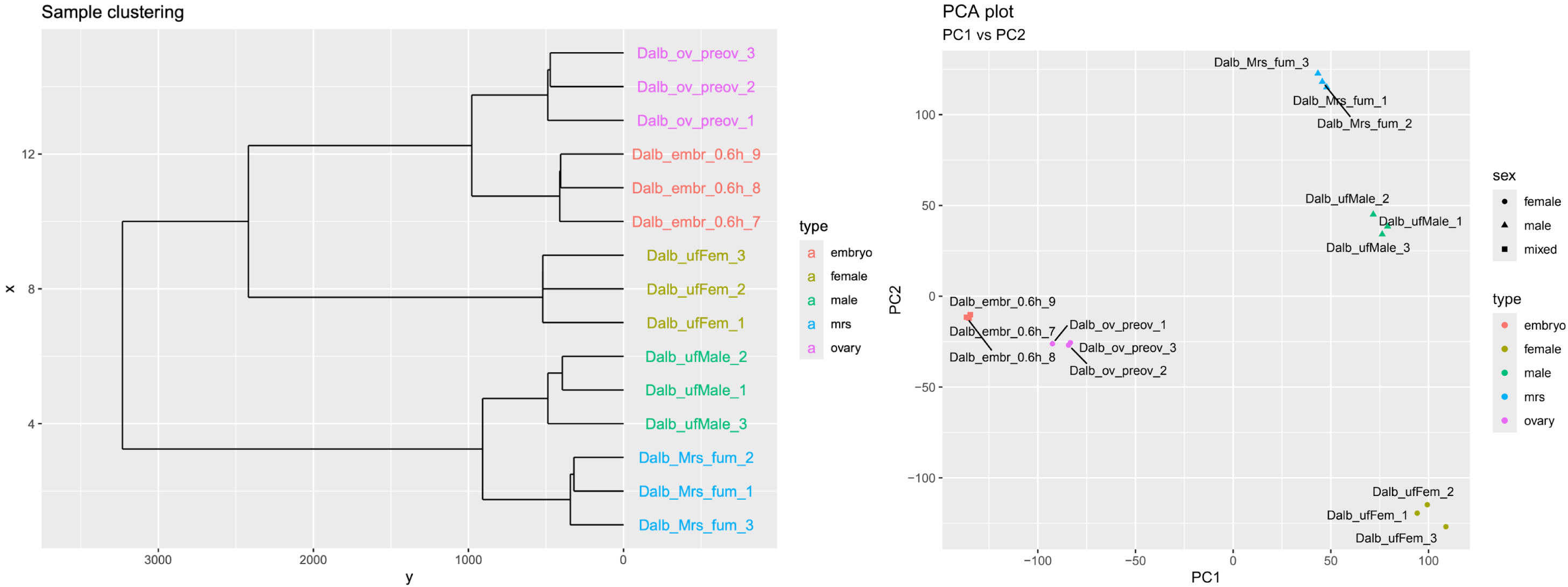
Clustering of *D.albipictus* RNA-seq samples comparing male, female and embryo sources of RNA. TPM values were used for clustering and PCA analyses. Dendrogram (A) and PCA analysis (B) showed the clustering of samples from the different sources of RNA.

**Table 3.** RNA sequencing FPKM data: FPKM_1_counts.xlsx.

| ID | FPKM > 1 count | FPKM > 1 mean | FPKM > 1 SD |
| --- | --- | --- | --- |
| Dalb_emb_0-6h_7 | 11809 |  |  |
| Dalb_emb_0-6h_8 | 11714 |  |  |
| Dalb_emb_0-6h_9 | 11766 | 11763 | 48 |
| Dalb_Mrs_fum_1 | 15025 |  |  |
| Dalb_Mrs_fum_2 | 14893 |  |  |
| Dalb_Mrs_fum_3 | 15040 | 14986 | 81 |
| Dalb_ov_preov_1 | 12037 |  |  |
| Dalb_ov_preov_2 | 11815 |  |  |
| Dalb_ov_preov_3 | 11794 | 11882 | 135 |
| Dalb_ufFem_1 | 11457 |  |  |
| Dalb_ufFem_2 | 11433 |  |  |
| Dalb_ufFem_3 | 11119 | 11336 | 189 |
| Dalb_ufMale_1 | 15878 |  |  |
| Dalb_ufMale_2 | 16099 |  |  |
| Dalb_ufMale_3 | 15524 | 15834 | 290 |

### Identification of Prevalent Functional Annotations and Gene Ontologies

Once transcripts were assembled, functional annotation was conducted using ORFfinder and HMMSCAN. A total of 11,933 Pfam domains were identified; 6,870 were identified two or more times and 5,063 were identified once within the data. Approximately 70% (15,377 of 22,680) of the genes had at least one Pfam domain, with 4,382 genes annotated with a single Pfam domain and 10,995 genes annotated with multiple Pfam domains. The top 20 most abundant Pfam domains consisted of 11 linked to zinc finger domains (PF00096, PF13894, PF13465, PF12874, PF12171, PF13912, PF12756, PF21816, PF00097, PF13920, PF13639), two linked to M13 peptidase (PF05649 and PF01431), two AAA+ ATPase domains (PF13401 and PF13191), in addition to a Protein kinase domain (PF00069), a Protein tyrosine and serine/threonine kinase (PF07714), a 50S ribosome-binding GTPase (PF01926), a Major Facilitator Superfamily (PF07690) and a WD domain (PF00400) (**Figure 3A**; **Table 4**).

**Figure 3.**
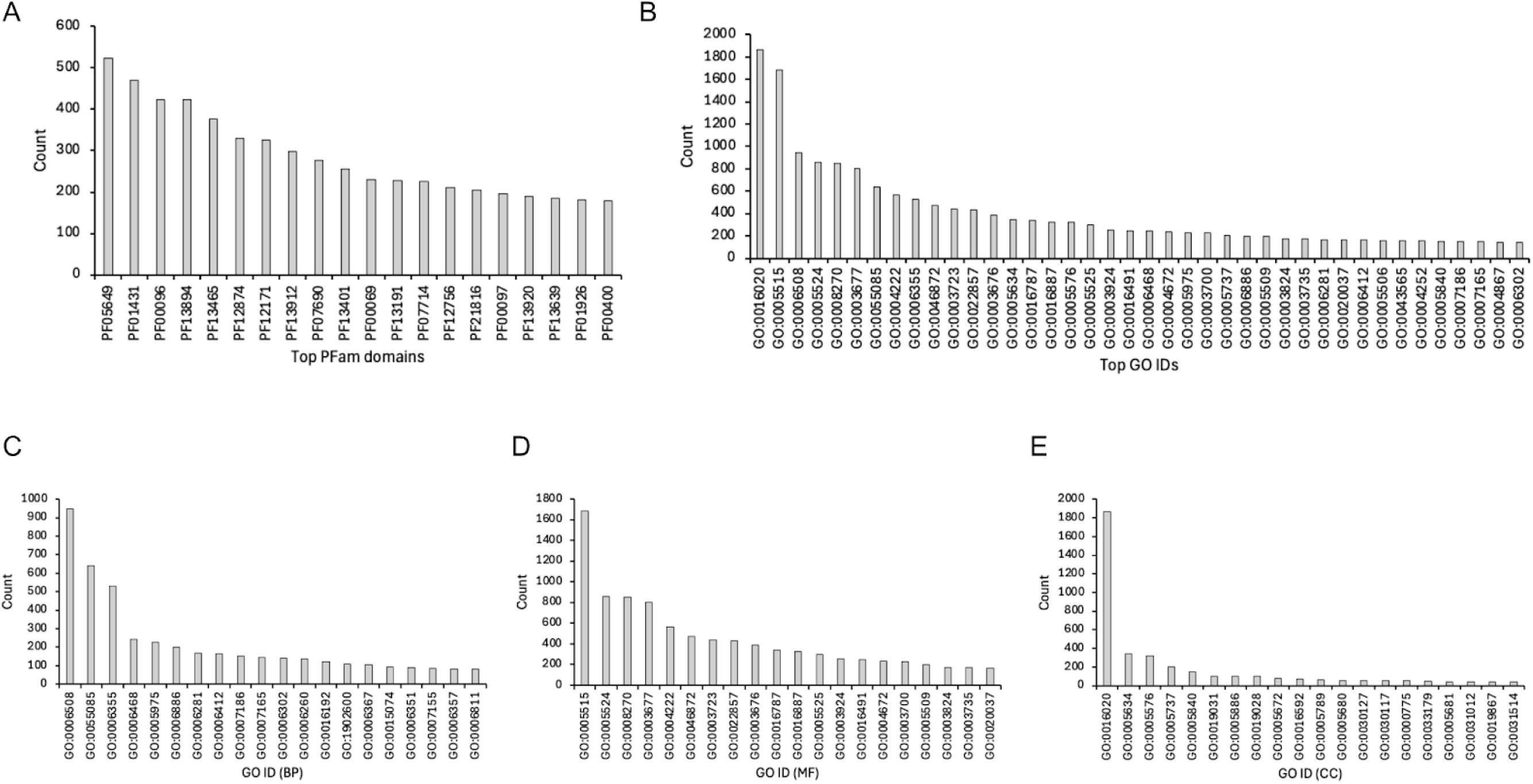
Identification of the most abundant Pfam domains and GO IDs assigned to sequences from *D.albipictus*. Transcripts were assembled and PFAM domain annotation was conducted using HMMSCAN (A). GO terms were assigned to genes based on PFAM-to-GO mapping. Top 40 most frequent terms are shown (B). The top 20 terms for each ontology (BP, MF, CC) were identified (C), (D), and (E) respectively.

**Table 4.** Top 20 most abundant PFAM domains identified in *D.albipictus*.

| *PFAM ID* | *PFAM Name* | *Total counts* |
| --- | --- | --- |
| PF05649 | Peptidase_M13_N | 524 |
| PF01431 | Peptidase_M13 | 470 |
| PF00096 | zf-C2H2 | 424 |
| PF13894 | zf-C2H2_4 | 423 |
| PF13465 | zf-H2C2_2 | 376 |
| PF12874 | zf-met | 330 |
| PF12171 | zf-C2H2_jaz | 326 |
| PF13912 | zf-C2H2_6 | 297 |
| PF07690 | MFS_1 | 276 |
| PF13401 | AAA_22 | 255 |
| PF00069 | Pkinase | 230 |
| PF13191 | AAA_16 | 227 |
| PF07714 | PK_Tyr_Ser-Thr | 225 |
| PF12756 | zf-C2H2_2 | 210 |
| PF21816 | Zap1_zf1 | 204 |
| PF00097 | zf-C3HC4 | 195 |
| PF13920 | zf-C3HC4_3 | 189 |
| PF13639 | zf-RING_2 | 186 |
| PF01926 | MMR_HSR1 | 180 |
| PF00400 | WD40 | 179 |

Genes (n=10,861) were annotated with Gene Ontology (GO) terms using PFAM-to-GO mapping with 7,733 genes associated with multiple terms. A total of 2,010 unique GO terms were assigned to the sequence data across all samples. The most numerous GO annotations were identified for sequences containing one or more annotations (**Figure 3B, 3C, 3D, 3E**; **Table 5**). These included GO:0016020 (membrane), GO:0005515 (protein binding), GO:0006508 (proteolysis), GO:0008270 (zinc ion binding), GO:0005524 (ATP binding), GO:0003677 (DNA binding), GO:0055085 (transmembrane transport), GO:0004222 (metalloendopeptidase activity), GO:0006355 (regulation of DNA-templated transcription), GO:0046872 (metal ion binding).

**Table 5.** Top 20 most abundant GO terms for biological process Molecular function and Cellular components from *D.albipictus*.

| *Biological process* |  |  | *Molecular function* |  |  | *Cellular component* |  |  |
| --- | --- | --- | --- | --- | --- | --- | --- | --- |
| *GO term* | *GO Name* | *Total counts* | *GO term* | *GO Name* | *Total counts* | *GO term* | *GO Name* | *Total counts* |
| GO:0006508 | proteolysis | 948 | GO:0005515 | protein binding | 1687 | GO:0016020 | membrane | 1864 |
| GO:0055085 | transmembrane transport | 640 | GO:0005524 | ATP binding | 861 | GO:0005634 | nucleus | 343 |
| GO:0006355 | regulation of DNA-templated transcription | 530 | GO:0008270 | zinc ion binding | 848 | GO:0005576 | extracellular region | 321 |
| GO:0006468 | protein phosphorylation | 243 | GO:0003677 | DNA binding | 805 | GO:0005737 | cytoplasm | 207 |
| GO:0005975 | carbohydrate metabolic process | 229 | GO:0004222 | metalloendopeptidase activity | 564 | GO:0005840 | ribosome | 152 |
| GO:0006886 | intracellular protein transport | 200 | GO:0046872 | metal ion binding | 474 | GO:0019031 | viral envelope | 103 |
| GO:0006281 | DNA repair | 168 | GO:0003723 | RNA binding | 438 | GO:0005886 | plasma membrane | 103 |
| GO:0006412 | translation | 164 | GO:0022857 | transmembrane transporter activity | 431 | GO:0019028 | viral capsid | 101 |
| GO:0007186 | G protein-coupled receptor signaling pathway | 151 | GO:0003676 | nucleic acid binding | 386 | GO:0005672 | transcription factor TFIIA complex | 77 |
| GO:0007165 | signal transduction | 146 | GO:0016787 | hydrolase activity | 339 | GO:0016592 | mediator complex | 69 |
| GO:0006302 | double-strand break repair | 139 | GO:0016887 | ATP hydrolysis activity | 322 | GO:0005789 | endoplasmic reticulum membrane | 64 |
| GO:0006260 | DNA replication | 138 | GO:0005525 | GTP binding | 298 | GO:0005680 | anaphase-promoting complex | 57 |
| GO:0016192 | vesicle-mediated transport | 121 | GO:0003924 | GTPase activity | 254 | GO:0030127 | COPII vesicle coat | 56 |
| GO:1902600 | proton transmembrane transport | 110 | GO:0016491 | oxidoreductase activity | 247 | GO:0030117 | membrane coat | 55 |
| GO:0006367 | transcription initiation at RNA polymerase II promoter | 105 | GO:0004672 | protein kinase activity | 237 | GO:0000775 | chromosome, centromeric region | 54 |
| GO:0015074 | DNA integration | 94 | GO:0003700 | DNA-binding<br>transcription factor<br>activity | 225 | GO:0033179 | proton-transporting<br>V-type ATPase, V0<br>domain | 47 |
| GO:0006351 | DNA-templated<br>transcription | 90 | GO:0005509 | calcium ion binding | 198 | GO:0005681 | spliceosomal complex | 41 |
| GO:0007155 | cell adhesion | 84 | GO:0003824 | catalytic activity | 172 | GO:0031012 | extracellular matrix | 41 |
| GO:0006357 | regulation<br>of transcription by<br>RNA polymerase II | 83 | GO:0003735 | structural<br>constituent of<br>ribosome | 171 | GO:0019867 | outer membrane | 41 |
| GO:0006811 | monoatomic ion<br>transport | 83 | GO:0020037 | heme binding | 166 | GO:0031514 | motile cilium | 40 |

To provide additional insight into gene function, closest homologs for predicted *D. albipictus* transcripts and proteins were identified in two annotated ticks from the *Dermacentor* genus, *Dermacentor andersoni* and *Dermacentor silvarum*. Transcripts encoded by 19,848 (87.5%) and 19,101 (84.2%) *D. albipictus* genes had significant blastn hits (e-value < 0.01) against transcriptomes of *D. andersoni* and *D. silvarum*, respectively. A total of 16,624 (73.3%) and 16,153 (71.2%) genes displayed significant homology with *D. andersoni* and *D. silvarum* on the protein level (**Table 6**).

**Table 6.** Identification of homologs for *D.albipictus* in closely related tick species (*Dermacentor andersoni*, and *Dermacentor silvarum*). Predicted transcripts and proteins from the *D. albipictus* were blast searched and significant hits were identified showing significant homology. [Dalb_finalv2_combined.pfam_blast (2).xlsx](https://docs.google.com/spreadsheets/d/18tPw2eNWpQRuRtbA9xqQV6QP8Z6GVx8m/edit?usp=drive_web&ouid=106471113672753704061&rtpof=true)

### Differential Gene Expression and GO Enrichment Analyses

Five pairwise comparisons were performed to identify differentially expressed genes between embr_0-6h and adults (i.e. ufMale + ufFem) (**Table 7**), ufMale and ufFem (**Table 8**), Mrs and ufMale (**Table 9)**, Mrs and Ov-preov (**Table 10**), Ov-preovand ufFem (**Table 11**; **Figure 4**). Expression levels of 17,155, 12,171, 12,788, 17,622 and 14,360 genes were significantly affected (padj < 0.05) between the five pairs of samples, respectively (**Table 12**). Up and down regulated gene sets identified in the comparisons were used to perform biological process GO enrichment analyses to present a high-level overview of the observed changes.

**Figure 4.**
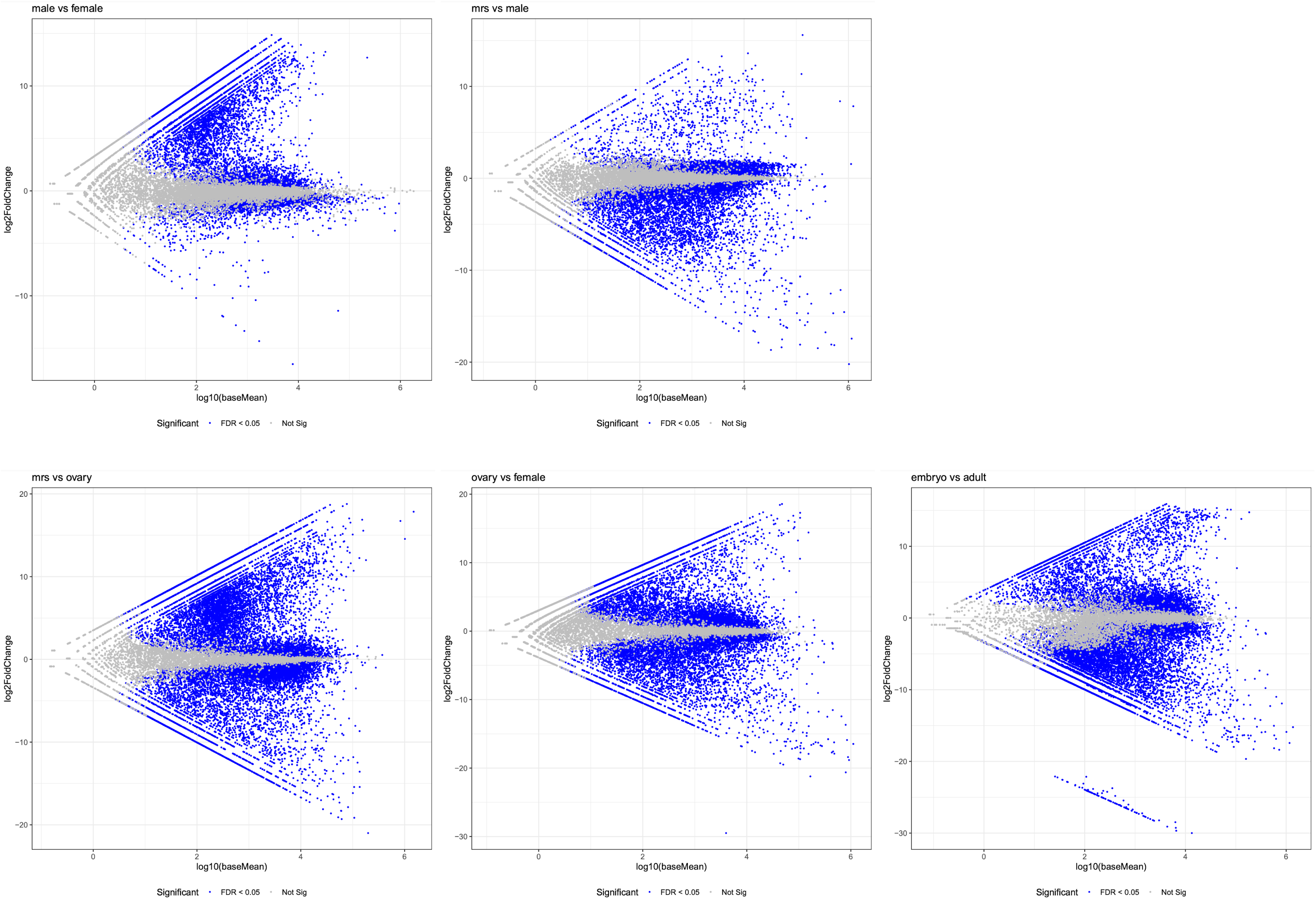
MA plots for pairwise analysis of differential gene expression in *D. albipictus* samples. Blue dots represent genes that are differentially expressed showing P-adjusted values <0.05. Gray dots represent genes that are not differentially expressed.

**Table 7.** Pairwise comparison using Deseq2 to identify differentially expressed genes between Embryos vs Adults. Data includes blast hits to *D.andersoni* and *D.silvarum* genomes. [embryo_vs_adult_deseq2.finalv2.pfam_blast.xlsx](https://docs.google.com/spreadsheets/d/1UOySBaBlm7_jLhovIGurgwfscry1ypt6/edit?gid=1604236441#gid=1604236441)

**Table 8.** Pairwise comparison using Deseq2 to identify differentially expressed genes between Male vs Female. Data includes blast hits to *D.andersoni* and *D.silvarum* genomes. [male_vs_female_deseq2.finalv2.pfam_blast.xlsx](https://docs.google.com/spreadsheets/d/1qaaIIc8lJcMSJGFk4gc8FcBFf01Vam7u/edit?gid=1339144774#gid=1339144774)

**Table 9.** Pairwise comparison using Deseq2 to identify differentially expressed genes between Mrs vs Male. Data includes blast hits to *D.andersoni* and *D.silvarum* genomes. [mrs_vs_male_deseq2.finalv2.pfam_blast.xlsx](https://docs.google.com/spreadsheets/d/1q8NG3iyvLXaSVaOkUvbd8LIPuXpJg54z/edit?usp=drive_web&ouid=106471113672753704061&rtpof=true)

**Table 10.** Pairwise comparison using Deseq2 to identify differentially expressed genes between Mrs vs Ovary. Data includes blast hits to *D.andersoni* and *D.silvarum* genomes. [mrs_vs_ovary_deseq2.finalv2.pfam_blast.xlsx](https://docs.google.com/spreadsheets/d/1vx-9OtXoPqp7w7gIytParKoLtNszuEYS/edit?usp=drive_web&ouid=106471113672753704061&rtpof=true)

**Table 11.** Pairwise comparison using Deseq2 to identify differentially expressed genes between Ovary vs Female. Data includes blast hits to *D.andersoni* and *D.silvarum* genomes. [ovary_vs_female_deseq2.finalv2.pfam_blast.xlsx](https://docs.google.com/spreadsheets/d/1NZmhiYxP2_pCMdK3n04yT3Oc1FnBbPS7/edit?gid=5945454#gid=5945454)

**Table 12.** Pairwise comparison of differential gene expression in *D.* albipictus showing statistically significant changes (padj <0.05) in gene expression and 20-fold change in expression.

| *ID* | *padj < 0.05*<br>*count* | *padj < 0.05 (%)* | *upregulated* | *downregulated* | *padj < 0.05 20*<br>*fold count* | *padj < 0.05 20*<br>*fold (%)* | *upregulated 20*<br>*fold* | *downregulated*<br>*20 fold* |
| --- | --- | --- | --- | --- | --- | --- | --- | --- |
| *embryo vs adult* | 17155 | 75.64 | 6895 | 10260 | 8702 | 38.37 | 2195 | 6507 |
| *male vs female* | 12171 | 53.66 | 8288 | 3883 | 5336 | 23.53 | 5242 | 94 |
| *mrs vs male* | 12788 | 56.38 | 5754 | 7034 | 2418 | 10.66 | 552 | 1866 |
| *mrs vs ovary* | 17622 | 77.7 | 9394 | 8228 | 7935 | 34.99 | 5458 | 2477 |
| *ovary vs female* | 14360 | 63.32 | 7936 | 6424 | 5329 | 23.5 | 3036 | 2293 |

For the embr_0-6h vs adult pair, top BP GO terms enriched in genes upregulated in embryos included GO:0006281 (DNA repair), GO:0006355 (regulation of DNA-templated transcription), GO:0006260 (DNA replication), GO:0000398 (mRNA splicing), GO:0000278 (mitotic cell cycle) (**Table 13**). Top terms enriched in adults were GO:0006508 (proteolysis), GO:0005975 (carbohydrate metabolic process), GO:0030682 (symbiont-mediated perturbation of host defenses), GO:0006412 (translation), GO:1900137 (negative regulation of chemokine activity) (**Table 13**). Consistent with this, some of the genes most strongly upregulated in embryos include MSTRG.18879, a putative homolog of BRISC and BRCA1-A complex member 1-like protein known to be involved in DNA repair-dependent chromatin remodeling and mitotic DNA damage checkpoint signaling. A number of genes encoding zinc finger containing proteins likely involved in various nucleic acid metabolic processes are also at the top of the embryonically enriched genes. On the other hand, a homolog of eukaryotic initiation factor 4A-I-like protein, MSTRG.2905, that is a part of the translation initiation complex, MSTRG.5657, which encodes a lysosomal alpha-mannosidase-like protein involved in glycoprotein break down, as well as a number of genes encoding various cuticle proteins are upregulated in adults (**Table 14**).

**Table 13.** Enriched biological process GO terms for embryo vs adult pairwise comparison. Top 20 up and down regulated terms were included from the data (embryo_vs_adult.topGO.fisher.bp.up.xlsx and embryo_vs_adult.topGO.fisher.bp.down.xlsx)

| Rank | upregulated |  |  |  |  |  | downregulated |  |  |  |  |  |
| --- | --- | --- | --- | --- | --- | --- | --- | --- | --- | --- | --- | --- |
|  | GO.ID | Term | Annotated | Significant | Expected | weight01fisher | GO.ID | Term | Annotated | Significant | Expected | weight01fisher |
| 1 | GO:0006281 | DNA repair | 341 | 175 | 114.65 | 1.20E-13 | GO:0006508 | proteolysis | 1139 | 573 | 474.99 | 1.00E-12 |
| 2 | GO:0006355 | regulation of DNA-templated transcription... | 651 | 306 | 218.87 | 1.20E-11 | GO:0005975 | carbohydrate metabolic process | 354 | 192 | 147.63 | 1.30E-08 |
| 3 | GO:0006260 | DNA replication | 207 | 108 | 69.59 | 2.20E-08 | GO:0030682 | symbiont-mediated perturbation of host d... | 63 | 48 | 26.27 | 2.40E-08 |
| 4 | GO:0000398 | mRNA splicing, via spliceosome | 95 | 57 | 31.94 | 1.00E-07 | GO:0006412 | translation | 253 | 134 | 105.51 | 1.90E-06 |
| 5 | GO:0000278 | mitotic cell cycle | 70 | 43 | 23.53 | 1.20E-07 | GO:1900137 | negative regulation of chemokine activit... | 15 | 14 | 6.26 | 4.30E-05 |
| 6 | GO:0006886 | intracellular protein transport | 229 | 114 | 76.99 | 2.00E-07 | GO:0055085 | transmembrane transport | 810 | 387 | 337.79 | 0.00013 |
| 7 | GO:0006888 | endoplasmic reticulum to Golgi vesicle-m... | 66 | 41 | 22.19 | 1.80E-06 | GO:0007596 | blood coagulation | 42 | 19 | 17.52 | 0.00092 |
| 8 | GO:0006351 | DNA-templated transcription | 865 | 402 | 290.82 | 4.30E-05 | GO:0006631 | fatty acid metabolic process | 48 | 28 | 20.02 | 0.00215 |
| 9 | GO:0016192 | vesicle-mediated transport | 257 | 131 | 86.4 | 4.80E-05 | GO:0016310 | phosphorylation | 254 | 92 | 105.92 | 0.00229 |
| 10 | GO:0017121 | plasma membrane phospholipid scrambling | 62 | 36 | 20.84 | 6.40E-05 | GO:2001256 | regulation of store-operated calcium ent... | 24 | 17 | 10.01 | 0.00369 |
| 11 | GO:0051604 | protein maturation | 201 | 85 | 67.58 | 7.90E-05 | GO:0036211 | protein modification process | 597 | 222 | 248.96 | 0.00456 |
| 12 | GO:0006352 | DNA-templated transcription initiation | 151 | 62 | 50.77 | 9.30E-05 | GO:0015986 | proton motive force-driven ATP synthesis | 23 | 16 | 9.59 | 0.0064 |
| 13 | GO:0006310 | DNA recombination | 155 | 76 | 52.11 | 0.00025 | GO:0009306 | protein secretion | 27 | 18 | 11.26 | 0.00766 |
| 14 | GO:0006364 | rRNA processing | 60 | 35 | 20.17 | 0.00036 | GO:0009166 | nucleotide catabolic process | 18 | 13 | 7.51 | 0.00815 |
| 15 | GO:0034080 | CENP-A containing chromatin assembly | 7 | 7 | 2.35 | 0.00048 | GO:0007218 | neuropeptide signaling pathway | 8 | 7 | 3.34 | 0.01111 |
| 16 | GO:0044237 | cellular metabolic process | 3016 | 1306 | 1014 | 0.00051 | GO:0046649 | lymphocyte activation | 5 | 5 | 2.09 | 0.01259 |
| 17 | GO:0007018 | microtubule-based movement | 43 | 25 | 14.46 | 0.00081 | GO:0019441 | tryptophan catabolic process to kynureni... | 5 | 5 | 2.09 | 0.01259 |
| 18 | GO:0000105 | histidine biosynthetic process | 18 | 13 | 6.05 | 0.00092 | GO:0036376 | sodium ion export across plasma membrane | 10 | 8 | 4.17 | 0.01631 |
| 19 | GO:0019051 | induction by virus of host apoptotic pro... | 18 | 13 | 6.05 | 0.00092 | GO:0050790 | regulation of catalytic activity | 62 | 29 | 25.86 | 0.02064 |
| 20 | GO:0015074 | DNA integration | 93 | 46 | 31.27 | 0.00107 | GO:0015031 | protein transport | 314 | 117 | 130.95 | 0.02216 |

**Table 14.** Differential gene expression for embryo vs adult *D.albipictus* samples. [embryo_vs_adult_deseq2.finalv2.pfam_blast.xlsx](https://docs.google.com/spreadsheets/d/1UOySBaBlm7_jLhovIGurgwfscry1ypt6/edit?gid=1604236441#gid=1604236441)

In ufMale vs ufFem comparison, GO:0006508 (proteolysis), GO:0005975 (carbohydrate metabolic process), GO:0030261 (chromosome condensation), GO:2001256 (regulation of store-operated calcium entry), GO:0006614 (SRP-dependent cotranslational protein targeting to membrane) terms were strongly enriched in genes upregulated in males (**Table 15**). In agreement with that, a number of peptidase-encoding genes, such as MSTRG.18259, MSTRG.10630, MSTRG.6630, MSTRG.3076 were among top male-biased genes, as were genes involved in carbohydrate metabolism including MSTRG.3527, MSTRG.12798, MSTRG.3172, the last one encoding a chitinase-3-like protein 1. Relatively few genes were upregulated in females and many of them did not have identifiable GO-annotated PFAM domains resulting in poor GO term enrichment with only GO:0006412 (translation) term being strongly significantly enriched. However, a several genes with homologs identified in *D. andersoni* and *D. silvarum* are significantly upregulated in females, including MSTRG.9157, which encodes an ETS homologous factor and transcription factor MafA-like gene MSTRG.16145 (**Table 16**).

**Table 15.** Enriched biological process GO terms for male vs female pairwise comparison. Top 20 up and down regulated terms were included from the data (male_vs_female.topGO.fisher.bp.up.xlsx and male_vs_female.topGO.fisher.bp.down.xlsx)

| upregulated |  |  |  |  |  |  | downregulated |  |  |  |  |  |
| --- | --- | --- | --- | --- | --- | --- | --- | --- | --- | --- | --- | --- |
| Rank | GO.ID | Term | Annotated | Significant | Expected | weight01fisher | GO.ID | Term | Annotated | Significant | Expected | weight01fisher |
| 1 | GO:0006508 | proteolysis | 1115 | 717 | 453.84 | < 1e-30 | GO:0006412 | translation | 245 | 96 | 50.77 | 2.30E-10 |
| 2 | GO:0005975 | carbohydrate metabolic process | 347 | 176 | 141.24 | 4.10E-08 | GO:0030431 | sleep | 8 | 8 | 1.66 | 3.30E-06 |
| 3 | GO:0030261 | chromosome condensation | 30 | 25 | 12.21 | 2.00E-06 | GO:0032222 | regulation of synaptic transmission, cho... | 8 | 8 | 1.66 | 3.30E-06 |
| 4 | GO:2001256 | regulation of store-operated calcium ent... | 24 | 19 | 9.77 | 0.00014 | GO:1903818 | positive regulation of voltage-gated pot... | 8 | 8 | 1.66 | 3.30E-06 |
| 5 | GO:0006614 | SRP-dependent cotranslational protein ta... | 64 | 40 | 26.05 | 0.00033 | GO:0007596 | blood coagulation | 40 | 11 | 8.29 | 0.0015 |
| 6 | GO:0006302 | double-strand break repair | 145 | 77 | 59.02 | 0.00056 | GO:0009058 | biosynthetic process | 2007 | 477 | 415.88 | 0.0015 |
| 7 | GO:0006270 | DNA replication initiation | 56 | 35 | 22.79 | 0.00078 | GO:0032508 | DNA duplex unwinding | 13 | 8 | 2.69 | 0.0016 |
| 8 | GO:0015031 | protein transport | 312 | 140 | 126.99 | 0.0012 | GO:0006813 | potassium ion transport | 58 | 26 | 12.02 | 0.0031 |
| 9 | GO:0016579 | protein deubiquitination | 33 | 22 | 13.43 | 0.00226 | GO:0006520 | amino acid metabolic process | 110 | 36 | 22.79 | 0.0032 |
| 10 | GO:0036211 | protein modification process | 578 | 245 | 235.26 | 0.0025 | GO:0007156 | homophilic cell adhesion via plasma memb... | 17 | 9 | 3.52 | 0.0033 |
| 11 | GO:1902600 | proton transmembrane transport | 110 | 59 | 44.77 | 0.00387 | GO:0055085 | transmembrane transport | 806 | 198 | 167.02 | 0.0037 |
| 12 | GO:0006397 | mRNA processing | 128 | 68 | 52.1 | 0.0065 | GO:0007186 | G protein-coupled receptor signaling pat... | 169 | 51 | 35.02 | 0.0047 |
| 13 | GO:0043666 | regulation of phosphoprotein phosphatase... | 8 | 7 | 3.26 | 0.0095 | GO:0050790 | regulation of catalytic activity | 62 | 19 | 12.85 | 0.0047 |
| 14 | GO:0019049 | virus-mediated perturbation of host defe... | 5 | 5 | 2.04 | 0.01115 | GO:0006633 | fatty acid biosynthetic process | 12 | 7 | 2.49 | 0.0048 |
| 15 | GO:0007059 | chromosome segregation | 92 | 46 | 37.45 | 0.02075 | GO:0007165 | signal transduction | 542 | 152 | 112.31 | 0.005 |
| 16 | GO:0002949 | tRNA threonylcarbamoyladenosine modifica... | 80 | 42 | 32.56 | 0.0211 | GO:0016042 | lipid catabolic process | 21 | 10 | 4.35 | 0.0053 |
| 17 | GO:0006511 | ubiquitin-dependent protein catabolic pr... | 99 | 53 | 40.3 | 0.02147 | GO:0006694 | steroid biosynthetic process | 40 | 15 | 8.29 | 0.0053 |
| 18 | GO:0009966 | regulation of signal transduction | 88 | 35 | 35.82 | 0.02488 | GO:0006418 | tRNA aminoacylation for protein translat... | 44 | 16 | 9.12 | 0.0063 |
| 19 | GO:0045892 | negative regulation of DNA-templated tra... | 10 | 7 | 4.07 | 0.06003 | GO:0030258 | lipid modification | 8 | 4 | 1.66 | 0.0089 |
| 20 | GO:0000398 | mRNA splicing, via spliceosome | 74 | 37 | 30.12 | 0.06536 | GO:0009152 | purine ribonucleotide biosynthetic proce... | 50 | 17 | 10.36 | 0.012 |

**Table 16.** Differential gene expression for male vs female *D.albipictus* samples. [male_vs_female_deseq2.finalv2.pfam_blast.xlsx](https://docs.google.com/spreadsheets/d/1qaaIIc8lJcMSJGFk4gc8FcBFf01Vam7u/edit?gid=1339144774#gid=1339144774)

The GO terms GO:0006508 (proteolysis), GO:0006302 (double-strand break repair), GO:0005975 (carbohydrate metabolic process), GO:0030261 (chromosome condensation), and GO:0007059 (chromosome segregation) were enriched in genes upregulated in Mrs compared to ufMale (**Table 17**) and peptidases such as MSTRG.10729, MSTRG.10218, MSTRG.20525, MSTRG.19268, MSTRG.16317 were among the strongest upregulated genes identified in this comparison (**Table 18**). GO:0030682 (symbiont-mediated perturbation of host defenses) is the most enriched GO term in genes overexpressed in ufMale and multiple proteins containing histamine binding domain, including MSTRG.4914 encoding a homolog of male-specific histamine-binding salivary protein are strongly expressed in male samples. These proteins are reported to function by sequestering histamine at the wound site thus suppressing inflammatory and immune responses of the host and enabling prolonged blood feeding by ticks. Other top ufMale enriched GO terms include GO:0006694 (steroid biosynthetic process), GO:0006811 (monoatomic ion transport), GO:0006811 (monoatomic ion transport), GO:1900137 (negative regulation of chemokine activity) (**Table 17**).

**Table 17.**
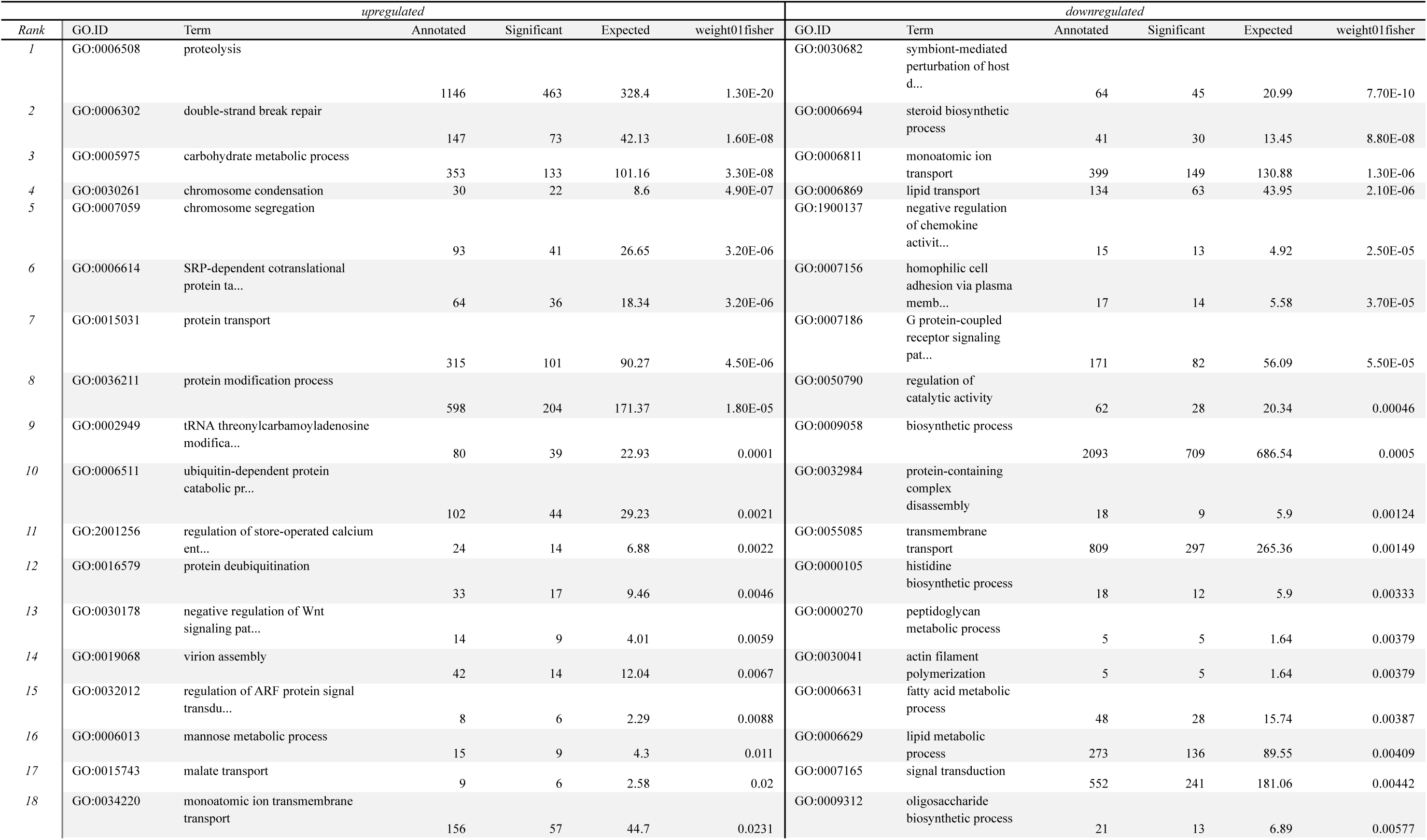
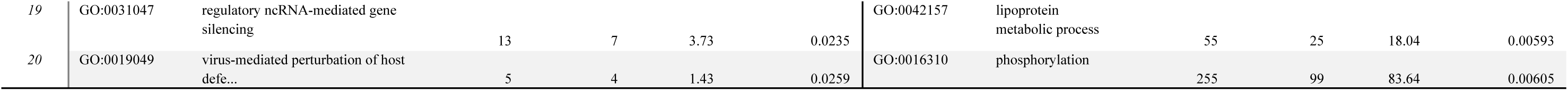
Enriched biological process GO terms for mrs vs male pairwise comparison. Top 20 up and down regulated terms were included from the data (mrs_vs_male.topGO.fisher.bp.up.xlsx and mrs_vs_male.topGO.fisher.bp.down.xlsx)

**Table 18.** Differential gene expression for mrs vs male *D.albipictus* samples. [mrs_vs_male_deseq2.finalv2.pfam_blast.xlsx](https://docs.google.com/spreadsheets/d/1q8NG3iyvLXaSVaOkUvbd8LIPuXpJg54z/edit?gid=649729190#gid=649729190)

Like in Mrs vs ufMale, Mrs vs Ov-preov comparison shows the enrichment of similar set of GO terms in Mrs samples, including GO:0006508 (proteolysis), GO:0005975 (carbohydrate metabolic process) and GO:0030261 (chromosome condensation) (**Table 19**). Genes expressed at higher levels in ovaries are enriched in GO:0015074 (DNA integration), GO:0006355 (regulation of DNA-templated transcription), GO:0006281 (DNA repair), GO:0006886 (intracellular protein transport), GO:0006801 (superoxide metabolic process). Genes annotated with these terms and are expressed at significantly higher levels in Ov-preov include ras-related protein Rab-32-like MSTRG.8423, forkhead box protein F1-like MSTRG.2103, coiled-coil domain-containing protein 22 homolog MSTRG.7996, K02A2.6-like MSTRG.5917, protein bicaudal C homolog 1-like MSTRG.824, zinc finger protein 425-like MSTRG.20876 (**Table 20**).

**Table 19.**
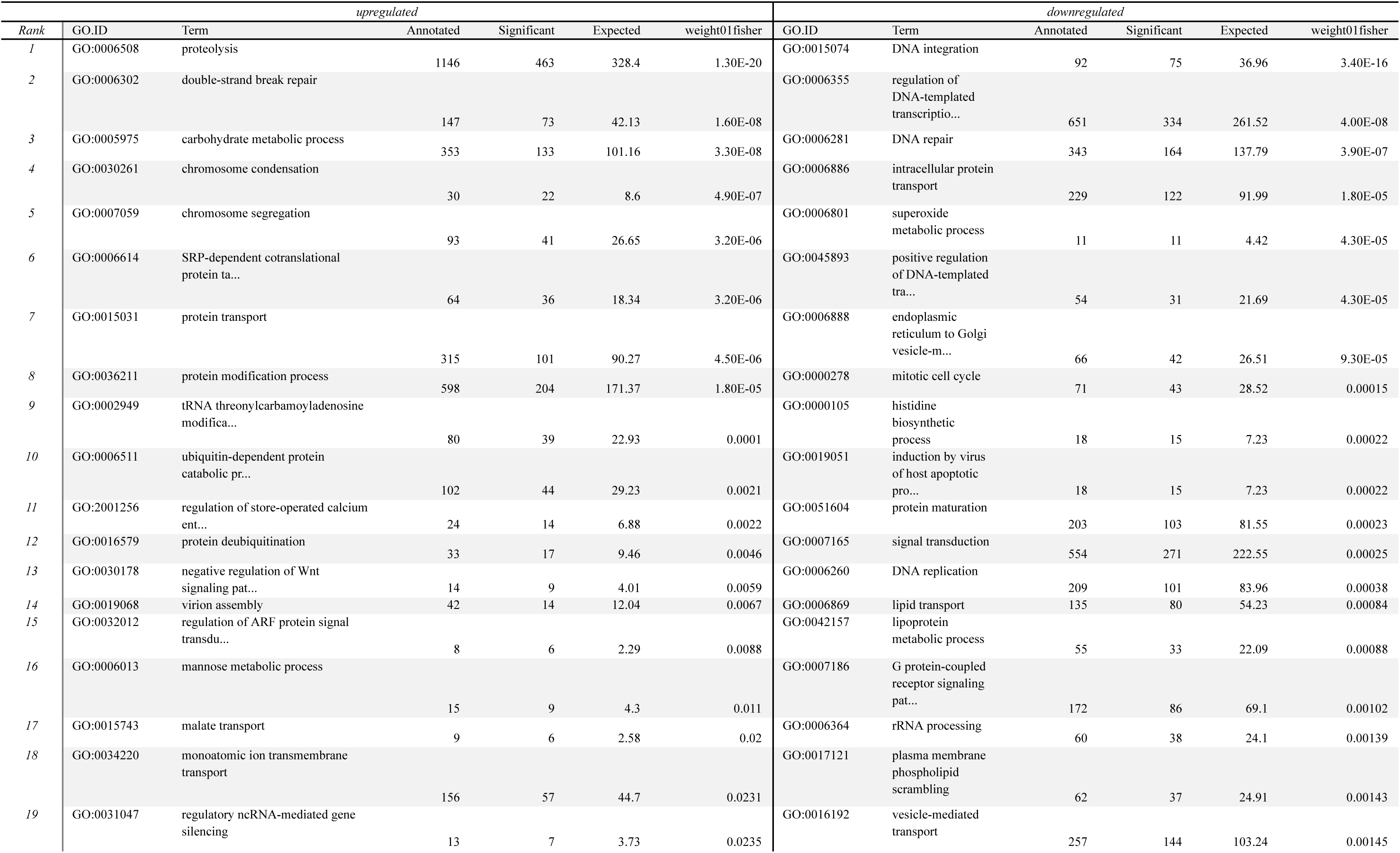
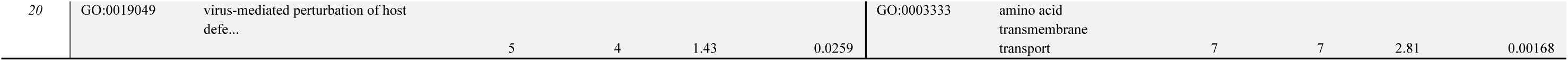
Enriched biological process GO terms for mrs vs ovary pairwise comparison. Top 20 up and down regulated terms were included from the data (mrs_vs_ovary.topGO.fisher.bp.up.xlsx and mrs_vs_ovary.topGO.fisher.bp.down.xlsx)

**Table 20.** Differential gene expression for mrs vs ovary *D.albipictus* samples. [mrs_vs_ovary_deseq2.finalv2.pfam_blast.xlsx](https://docs.google.com/spreadsheets/d/1vx-9OtXoPqp7w7gIytParKoLtNszuEYS/edit?gid=1906383320#gid=1906383320)

The ovary enriched set in the Ov-preov vs ufFem comparison resembles the one above and include such terms as GO:0015074 (DNA integration), GO:0006886 (intracellular protein transport), GO:0006260 (DNA replication), GO:0016192 (vesicle-mediated transport), GO:1902600 (proton transmembrane transport) (**Table 21**). Like in ufMale, GO:0030682 (symbiont-mediated perturbation of host defenses) is strongly enriched in ufFem and many histamine binding domain encoding genes that are likely involved in the suppression of the host immune response during the blood meal are overexpressed in females (MSTRG.19815, MSTRG.19824, MSTRG.4914, MSTRG.4465). 12 out of 15 genes annotated with GO:1900137 (negative regulation of chemokine activity) in the genome are strongly overexpressed in females compared to ovaries; MSTRG.6585, MSTRG.17014, MSTRG.6479, and MSTRG.6481 for example. All of these genes encode proteins containing the EVA_Class_A domain found in evasins proteins from ticks that have anti-inflammatory activity due to their binding to chemokines, which prevents leukocyte recruitment and increases host blood flow during feeding. Genes encoding Inhibitor_I68 domain containing proteins and annotated with GO:0007596 (blood coagulation) are also strongly overexpressed in females. These tick proteins are reported to be involved in the maintenance of the host blood liquidity during feeding. Many genes encoding structural components of ribosome and other translation related genes are also at the top of overexpressed gene list, e. g. MSTRG.16750, MSTRG.18888, MSTRG.19229 (**Table 21**; **Table 22**).

**Table 21.** Enriched biological process GO terms for ovary vs female pairwise comparison. Top 20 up and down regulated terms were included from the data (ovary_vs_female.topGO.fisher.bp.up.xlsx and ovary_vs_female.topGO.fisher.bp.down.xlsx)

| Rank | upregulated |  |  |  |  |  | downregulated |  |  |  |  |  |
| --- | --- | --- | --- | --- | --- | --- | --- | --- | --- | --- | --- | --- |
|  | GO.ID | Term | Annotated | Significant | Expected | weight01fisher | GO.ID | Term | Annotated | Significant | Expected | weight01fisher |
| 1 | GO:0015074 | DNA integration | 91 | 70 | 38.26 | 9.70E-12 | GO:0006412 | translation | 249 | 116 | 75.66 | 4.70E-10 |
| 2 | GO:0006886 | intracellular protein transport | 222 | 127 | 93.33 | 9.50E-07 | GO:0030682 | symbiont-mediated perturbation of host d... | 57 | 37 | 17.32 | 7.00E-08 |
| 3 | GO:0006260 | DNA replication | 202 | 113 | 84.92 | 1.50E-05 | GO:1900137 | negative regulation of chemokine activit... | 13 | 12 | 3.95 | 5.70E-06 |
| 4 | GO:0016192 | vesicle-mediated transport | 252 | 144 | 105.94 | 1.50E-05 | GO:0006811 | monoatomic ion transport | 383 | 128 | 116.37 | 7.10E-05 |
| 5 | GO:1902600 | proton transmembrane transport | 104 | 63 | 43.72 | 9.50E-05 | GO:0007596 | blood coagulation | 40 | 17 | 12.15 | 7.20E-05 |
| 6 | GO:0006364 | rRNA processing | 58 | 40 | 24.38 | 0.00016 | GO:0006694 | steroid biosynthetic process | 40 | 24 | 12.15 | 0.00014 |
| 7 | GO:0000278 | mitotic cell cycle | 70 | 47 | 29.43 | 0.0002 | GO:0055085 | transmembrane transport | 765 | 268 | 232.44 | 0.00021 |
| 8 | GO:0006281 | DNA repair | 338 | 183 | 142.09 | 0.00051 | GO:0030150 | protein import into mitochondrial matrix | 5 | 5 | 1.52 | 0.00258 |
| 9 | GO:0045893 | positive regulation of DNA-templated tra... | 54 | 29 | 22.7 | 0.00137 | GO:0016310 | phosphorylation | 248 | 82 | 75.35 | 0.00402 |
| 10 | GO:0034080 | CENP-A containing chromatin assembly | 7 | 7 | 2.94 | 0.00231 | GO:0006629 | lipid metabolic process | 261 | 109 | 79.3 | 0.00569 |
| 11 | GO:0030001 | metal ion transport | 179 | 73 | 75.25 | 0.00235 | GO:0010951 | negative regulation of endopeptidase act... | 12 | 8 | 3.65 | 0.01025 |
| 12 | GO:0006397 | mRNA processing | 149 | 87 | 62.64 | 0.00288 | GO:0006631 | fatty acid metabolic process | 48 | 22 | 14.58 | 0.01047 |
| 13 | GO:0006298 | mismatch repair | 15 | 12 | 6.31 | 0.00317 | GO:0006122 | mitochondrial electron transport, ubiqui... | 6 | 5 | 1.82 | 0.01157 |
| 14 | GO:0007059 | chromosome segregation | 93 | 50 | 39.1 | 0.0047 | GO:0007304 | chorion-containing eggshell formation | 8 | 6 | 2.43 | 0.01203 |
| 15 | GO:0000398 | mRNA splicing, via spliceosome | 94 | 52 | 39.52 | 0.00612 | GO:0015986 | proton motive force-driven ATP synthesis | 22 | 12 | 6.68 | 0.01544 |
| 16 | GO:0051301 | cell division | 71 | 37 | 29.85 | 0.007 | GO:0019915 | lipid storage | 15 | 9 | 4.56 | 0.01652 |
| 17 | GO:0006888 | endoplasmic reticulum to Golgi vesicle-m... | 66 | 38 | 27.75 | 0.00759 | GO:0007186 | G protein-coupled receptor signaling pat... | 169 | 64 | 51.35 | 0.02652 |
| 18 | GO:0000413 | protein<br>peptidyl-prolyl<br>isomerization | 11 | 9 | 4.62 | 0.0087 | GO:0043093 | FtsZ-dependent<br>cytokinesis | 39 | 18 | 11.85 | 0.02711 |
| 19 | GO:0031109 | microtubule<br>polymerization or<br>depolymeri... | 5 | 5 | 2.1 | 0.0131 | GO:0090529 | cell septum assembly | 39 | 18 | 11.85 | 0.02711 |
| 20 | GO:0000462 | maturation of<br>SSU-rRNA from<br>tricistronic... | 5 | 5 | 2.1 | 0.0131 | GO:0008610 | lipid biosynthetic<br>process | 126 | 53 | 38.28 | 0.02746 |

**Table 22.** Differential gene expression for ovary vs female *D.albipictus* samples. [ovary_vs_female_deseq2.finalv2.pfam_blast.xlsx](https://docs.google.com/spreadsheets/d/1NZmhiYxP2_pCMdK3n04yT3Oc1FnBbPS7/edit?gid=5945454#gid=5945454)

### Identification of *Doublesex* as a Target for Genetic Control Tools

In many insect species, the *doublesex* gene acts as a major sex determination factor and has been used for engineering of genetic control tools (Kyrou et al. 2018; Kandul et al. 2019; Li et al. 2024). Our blast searches identified four genes that encode proteins homologous to doublesex-like proteins in *D. andersoni* and *D. silvarum,* MSTRG.10335, MSTRG.10337, MSTRG.18500 and MSTRG.9862. MSTRG.10335 and MSTRG.10337 are located adjacent to each other on scaffold CM090517.1, MSTRG.9862 is also on CM090517.1 more than 50 Mb away from the MSTGR.10335/ MSTGR.10337 locus and MSTRG.18500 - on scaffold CM090522.1. MSTRG.10337 was identified as one of the strongly differentially expressed genes in the male vs female comparison. It is practically absent in female samples but is expressed at TPM of ∼9 in males (**Table 23**). The strong male-specific expression makes MSTRG.10337 a good candidate for possible sex determination regulator, which can be tested by inducing its expression in females or suppressing it in males. MSTRG.10335 is robustly expressed in both male and female samples with slightly higher, although not significantly, levels in males. However, it displays a profound difference in splicing pattern between ufMale and ufFem as well as between the germline samples Mrs and Ov-Preov. Splice junction SJ004 appears to be female specific, while SJ005 and SJ006 – male specific (**Table 24**). SJ004 belongs to MSTRG.10335.3 transcript, which is exclusive to ufFem and Ov-Preov samples, where it is expressed at the highest levels. SJ006 belongs to the MSTRG.10335.1 transcript, which is ufMale and Mrs specific. SJ005 originates from MSTRG.10335.2, which is not expressed in Ov-Preov and is expressed at very low levels in ufFem samples. Since sex-specific splicing of *doublesex* is necessary for proper sex determination in multiple species, including model organisms *Drosophila melanogaster* and *Bombyx mori* (Suzuki et al. 2001), and mosquitoes *A. aegypti* and *A. albopictus* (Salvemini et al. 2011; Jin et al. 2021), the splicing pattern of MSTRG.10335 could be explored for engineering strains for population control via pgSIT or for fluorescent marker-based sex sorting. MSTRG.9862 is expressed strongly at ∼10-12 TPM levels in both males and females and displays no differential splicing. Likewise, MSTRG.18500 shows no differential expression or splicing between male and female samples and is also expressed at low levels in all samples except for embr_0-6h. This makes MSTRG.9862 and MSTRG.18500 less likely to be involved in sex determination, although additional studies are required for confirmation. Furthermore, additional *doublesex* candidates may be discovered as more RNA-seq data are accumulated and better annotations become available.

**Table 23.** Expression of four putative *doublesex* genes identified in *D.albipictus* sequencing data. [dsx_candidates_expression.xlsx](https://docs.google.com/spreadsheets/d/1op_wv0EZ9TlxYRDpXdRfjOKmNyrOn9-n/edit?gid=866635258#gid=866635258)

**Table 24.** Splice junction analysis of MSTRG 10335 showing the sex specificity of SJ004 (female, green) and SJ005 and SJ006 (male, blue).

| Junction ID | Mrs_fum<br>1 | Mrs_fum<br>2 | Mrs_fum<br>3 | ov_preov<br>1 | ov_preov<br>2 | ov_preov<br>3 | ufFem<br>1 | ufFem<br>2 | ufFem<br>3 | ufMale<br>1 | ufMale<br>2 | ufMale<br>3 | Mrs_me<br>an | ov_me<br>an | ufFem_me<br>an | ufMale_me<br>an |
| --- | --- | --- | --- | --- | --- | --- | --- | --- | --- | --- | --- | --- | --- | --- | --- | --- |
| MSTRG.10335:SJ<br>001 | 19 | 18 | 15 | 4 | 5 | 10 | 5 | 7 | 0 | 18 | 22 | 12 | 17.33 | 12.00 | 4.00 | 17.33 |
| MSTRG.10335:SJ<br>002 | 23 | 24 | 17 | 2 | 5 | 10 | 3 | 0 | 1 | 13 | 25 | 27 | 21.33 | 8.00 | 1.33 | 21.67 |
| MSTRG.10335:SJ<br>003 | 20 | 22 | 9 | 1 | 3 | 3 | 2 | 6 | 2 | 19 | 23 | 19 | 17.00 | 6.67 | 3.33 | 20.33 |
| MSTRG.10335:SJ<br>004 | 0 | 0 | 0 | 3 | 2 | 8 | 1 | 1 | 1 | 0 | 0 | 0 | 0.00 | 4.33 | 1.00 | 0.00 |
| MSTRG.10335:SJ<br>005 | 5 | 12 | 6 | 0 | 0 | 0 | 1 | 1 | 0 | 5 | 14 | 7 | 7.67 | 0.00 | 0.67 | 8.67 |
| MSTRG.10335:SJ<br>006 | 8 | 6 | 2 | 0 | 0 | 0 | 0 | 0 | 0 | 15 | 7 | 18 | 5.33 | 0.00 | 0.00 | 13.33 |

### Analysis of Global Sex-Specific Splicing Patterns

To identify potential additional targets for splicing-based strain engineering, we performed exon utilization analysis between male and female samples on the whole genome level. Among 19,219 genes with multiple transcripts, 6,852 showed evidence of sex-specific splicing (padj < 0.1), affecting 26,814 out of 190,156 exons (**Table 25**). Exon utilization analysis identifies differentially expressed exons while controlling for differences due to overall gene expression levels in the samples. However, it cannot fully reconstruct alternatively spliced transcript structures and is highly sensitive to the quality of genomic annotations. Therefore, manual examination of transcript structures and read alignments is necessary to confirm the validity of identified hits. While the examination of the top candidates generated by the analysis identified several false positives due to incorrect gene annotations (e.g. Chimeric transcripts), multiple true sex-specific splicing events were identified.

**Table 25.** Exon utilization Analysis in D. albipictus male vs female pairwise comparison. [male_vs_female.dexseq.finalv2.xlsx](https://docs.google.com/spreadsheets/d/1w_kr9ftM5T_YFUktU6dCwzjZZaiH1f1R/edit?gid=641968034#gid=641968034)

One example, MSTRG.4611, encodes a homolog of a fatty acid CoA ligase Acsl3-like protein and has six annotated transcripts (**Figure 5**). Exonic parts E004-6 and E012 were significantly downregulated in male samples (padj < 1e-55; log2 fold change > 8) (**Figure 6**). Splice junction counts confirmed male specificity of several junctions: SJ002 connects the first two exons in isoforms MSTRG.4611.4 and MSTRG.4611.6, while SJ003 is specific to MSTRG.4611.3-6. Additionally, SJ008 and SJ010 splice exonic part E012, which is present in transcripts MSTRG.46112.2-4 (**Table 26**). These findings indicate that four transcripts with transcription start sites (TSS) near CM090514.1:436734180 are male specific, while two transcripts with TSS near CM090514.1:436713550 are expressed in both sexes. Protein alignment revealed that the male-specific exon E012 inserts an MLPV amino acid sequence into the active site of the eukaryotic long-chain fatty acid CoA synthetase domain (LC-FACS). This modification may alter the substrate specificity of male isoforms (**Figure 7**).

**Figure 5.**
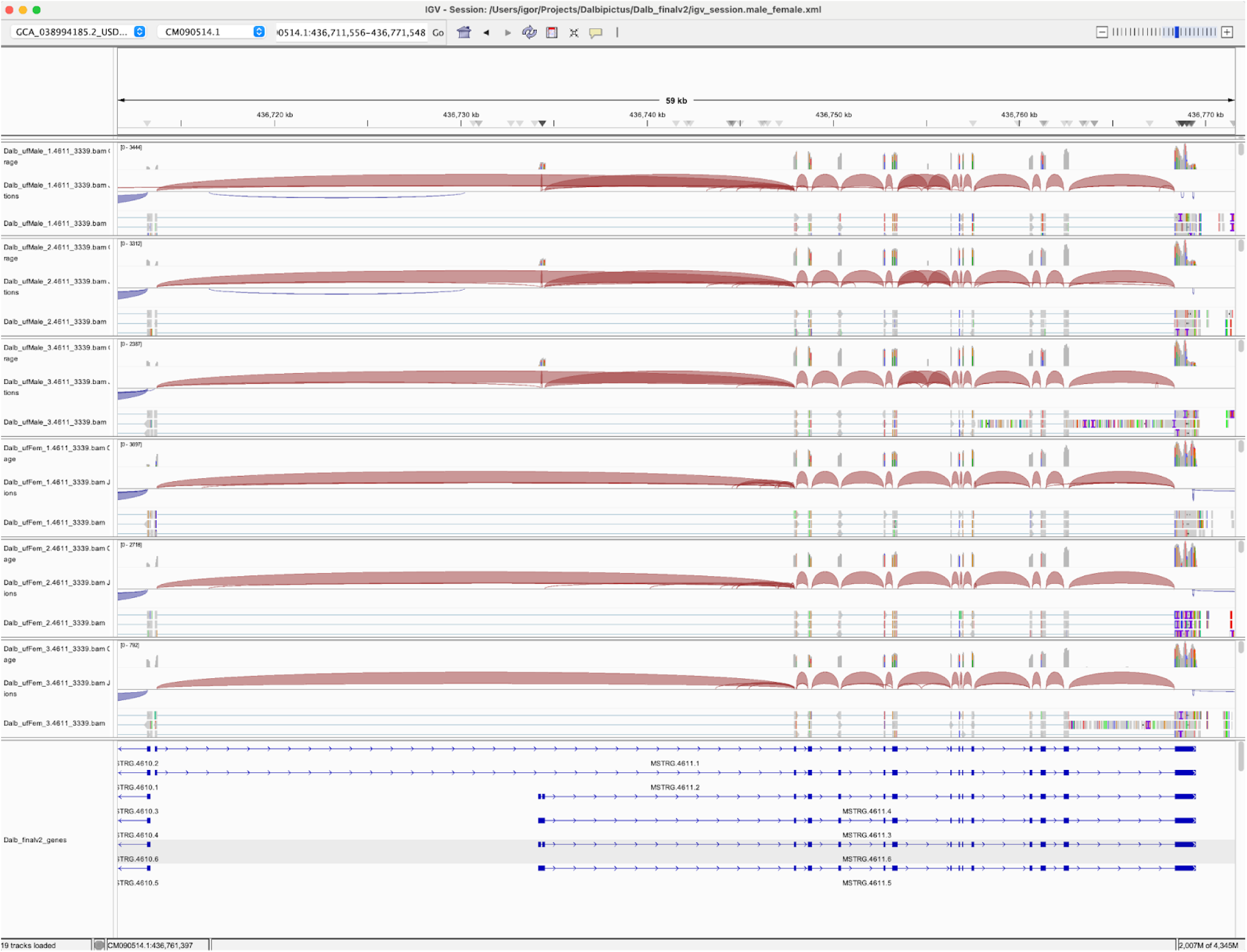
IGV transcript plot comparing male vs female *D.albipictus* splicing patterns for MSTRG 4611.

**Figure 6.**
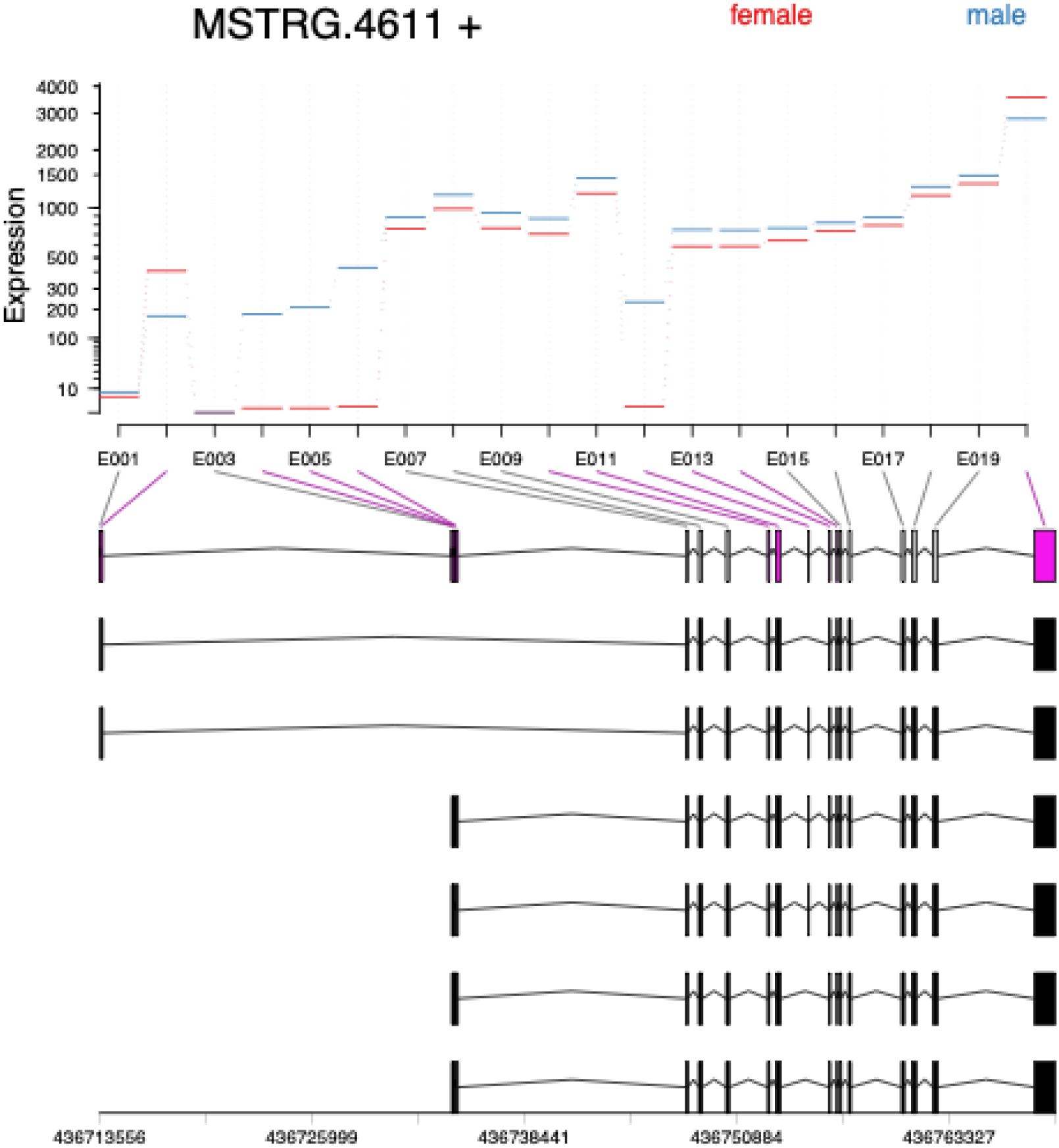
Exon plots analysis for MSTRG 4611 to elucidate sex specificity in *D. albipictus.* Exons were quantified, followed by differential usage analysis for the three ufMale compared to the three ufFem samples. Exons E004-6 and E012 show downregulation in male samples.

**Figure 7.**
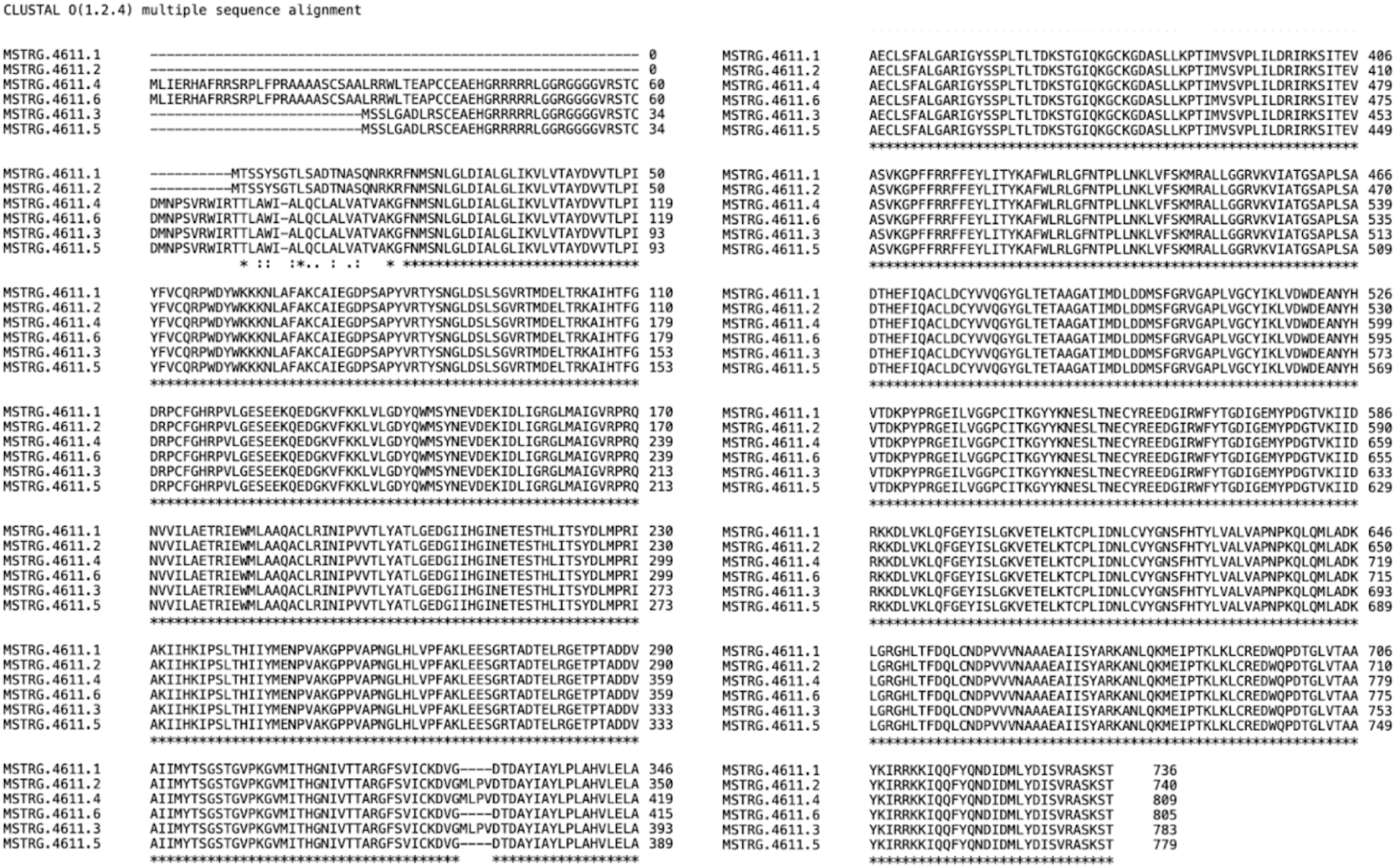
Protein alignment of MSTRG4611 transcripts MSTRG.4611.1-6 using Clustal. MLVP amino acid sequence is inserted into the male transcripts MSTRG.4611.2-4 in exon 12.

**Table 26.**
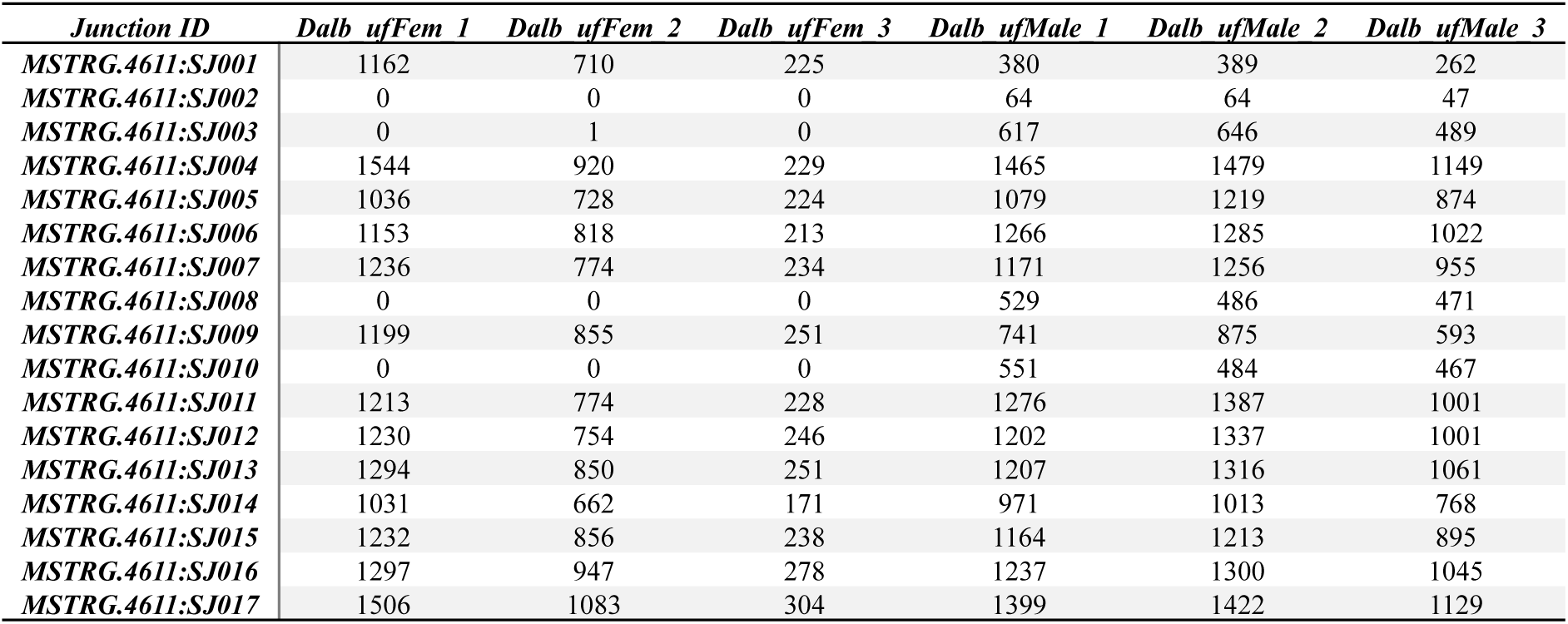
Splice junction counts for D. albipictus MSTRG 4611 showing sex-specificity of SJ002, SJ003, SJ008, and SJ010 (male).

| *Junction ID* | *Dalb ufFem 1* | *Dalb ufFem 2* | *Dalb ufFem 3* | *Dalb ufMale 1* | *Dalb ufMale 2* | *Dalb ufMale 3* |
| --- | --- | --- | --- | --- | --- | --- |
| *MSTRG.4611: SJ001* | 1162 | 710 | 225 | 380 | 389 | 262 |
| *MSTRG.4611: SJ002* | 0 | 0 | 0 | 64 | 64 | 47 |
| *MSTRG.4611: SJ003* | 0 | 1 | 0 | 617 | 646 | 489 |
| *MSTRG.4611: SJ004* | 1544 | 920 | 229 | 1465 | 1479 | 1149 |
| *MSTRG.4611: SJ005* | 1036 | 728 | 224 | 1079 | 1219 | 874 |
| *MSTRG.4611: SJ006* | 1153 | 818 | 213 | 1266 | 1285 | 1022 |
| *MSTRG.4611: SJ007* | 1236 | 774 | 234 | 1171 | 1256 | 955 |
| *MSTRG.4611: SJ008* | 0 | 0 | 0 | 529 | 486 | 471 |
| *MSTRG.4611: SJ009* | 1199 | 855 | 251 | 741 | 875 | 593 |
| *MSTRG.4611: SJ010* | 0 | 0 | 0 | 551 | 484 | 467 |
| *MSTRG.4611: SJ011* | 1213 | 774 | 228 | 1276 | 1387 | 1001 |
| *MSTRG.4611: SJ012* | 1230 | 754 | 246 | 1202 | 1337 | 1001 |
| *MSTRG.4611: SJ013* | 1294 | 850 | 251 | 1207 | 1316 | 1061 |
| *MSTRG.4611: SJ014* | 1031 | 662 | 171 | 971 | 1013 | 768 |
| *MSTRG.4611: SJ015* | 1232 | 856 | 238 | 1164 | 1213 | 895 |
| *MSTRG.4611: SJ016* | 1297 | 947 | 278 | 1237 | 1300 | 1045 |
| *MSTRG.4611: SJ017* | 1506 | 1083 | 304 | 1399 | 1422 | 1129 |

Another example is MSTRG.10562, which encodes ralA-binding protein 1-like involved in small GTPase-mediated signal transduction and comprises three transcripts (**Figure 8**). Exon 7 was significantly downregulated in males (padj =2.4e-145;log2 fold change =2.91) (**Figure 9**). Splice junction counts showed that SJ006, which connects exons 6 and 8, is male-specific, while SJ005 and SJ007 are enriched in females (**Table 27**). These results confirm that isoforms MSTRG.10562.1 and MSTRG.10562.3 are exclusively expressed in males. Protein alignment indicated that male-specific isoforms lack a 125 amino acid stretch in the second half of the protein (**Figure 10**). Although no conserved domains were identified in this region, it may play a role in modulating the efficiency of the GTPase activation cascade.

**Figure 8.**
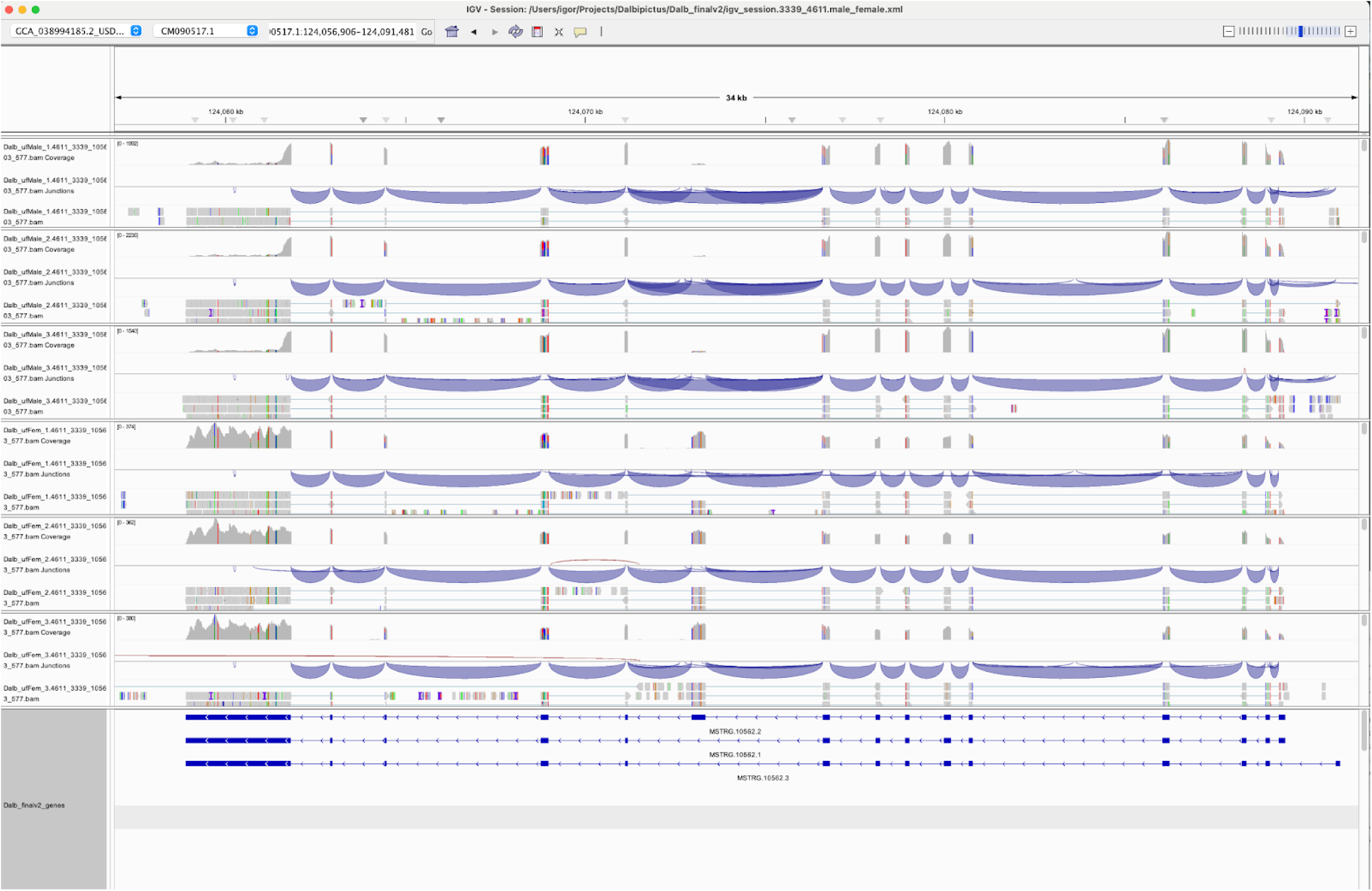
IGV transcript plot comparing male vs female *D.albipictus* splicing patterns for MSTRG 15062.

**Figure 9.**
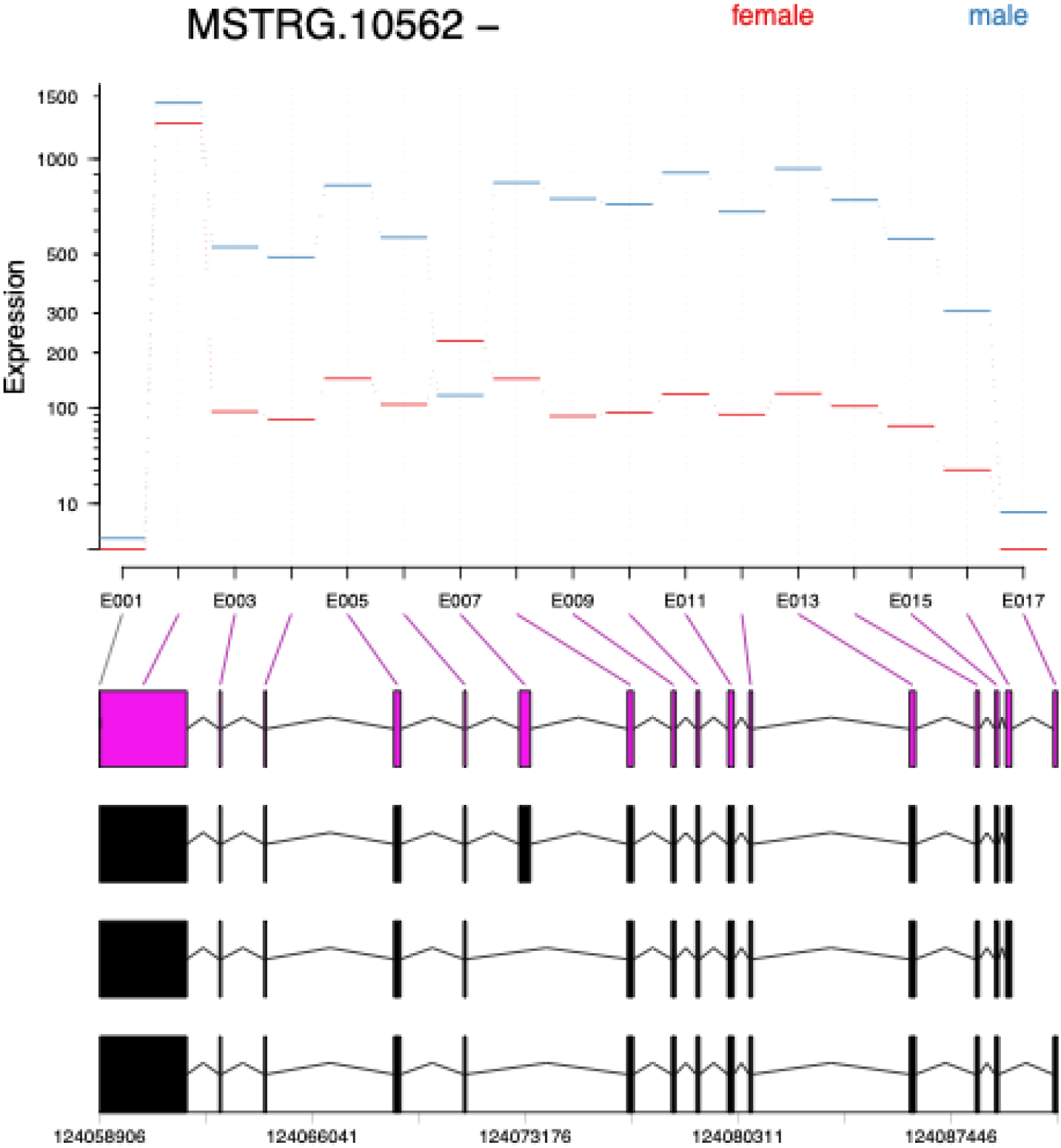
Exon plots analysis for MSTRG 10562 to elucidate sex specificity in *D. albipictus.* Exons were quantified, followed by differential usage analysis for the three ufMale compared to the three ufFem samples. Exons E007 show downregulation in male samples.

**Figure 10.**
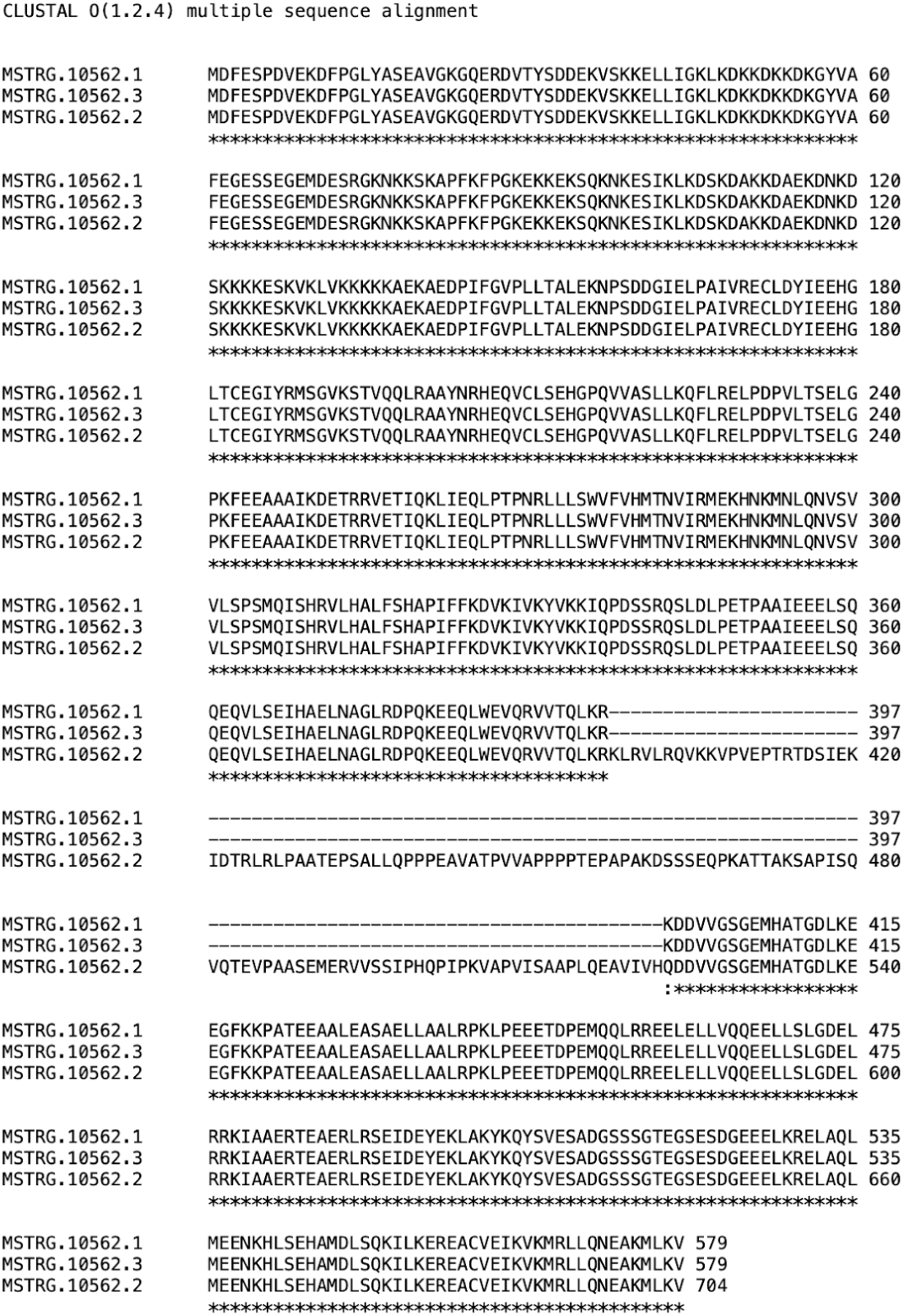
Protein alignment of MSTRG 10562 transcripts MSTRG.10562.1-3 using Clustal. 125 amino acid sequence is absent in the male transcripts MSTRG.10562.1-2.

**Table 27.**
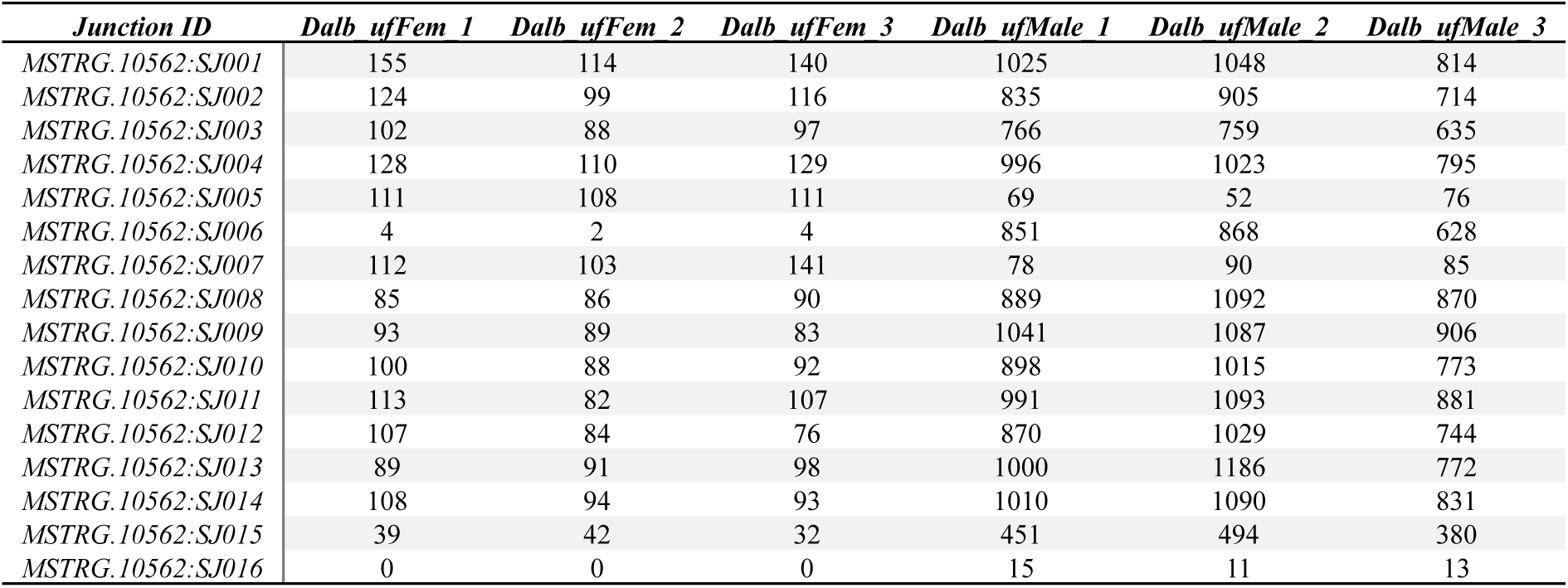
Splice junction counts for D. albipictus MSTRG 10562 showing sex-specificity of SJ006 (male), and SJ005, SJ007 (female enriched).

A third example is MSTRG.3339, which encodes a homolog of aquaporin-9 and expresses four isoforms (**Figure 11**). Exonic parts E009 and E010, along with corresponding splice junctions SJ006, SJ009, and SJ010, were exclusively expressed in male samples and are associated with isoforms MSTRG.3339.3 and MSTRG.3339.4 (**Figure 12**; **Table 28**). Alternative splicing alters the carboxyl terminus of the encoded proteins, which lies just outside the major intrinsic protein (MIP) domain predicted to form a membrane channel (**Figure 13**).

**Figure 11.**
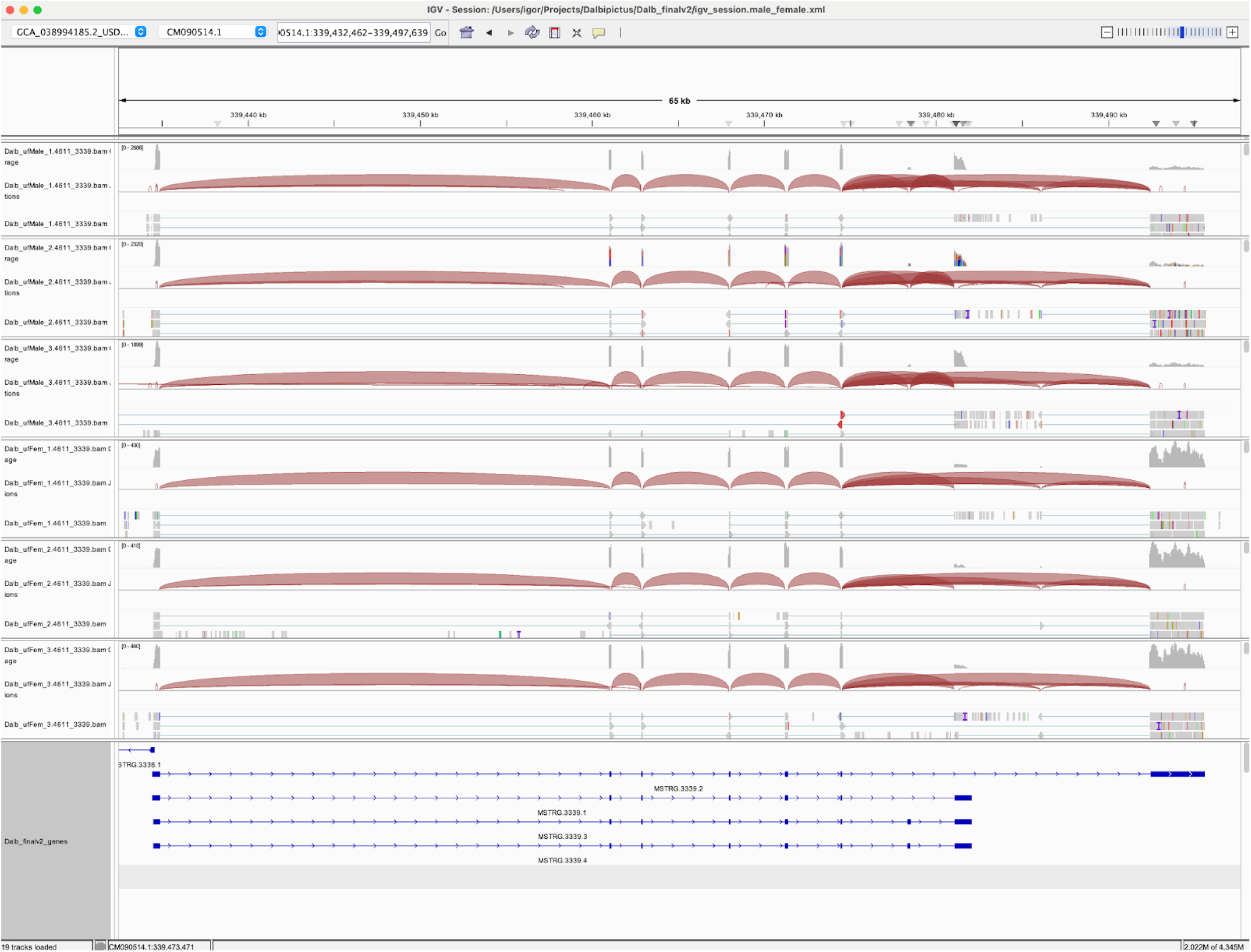
IGV transcript plot comparing male vs female *D.albipictus* splicing patterns for MSTRG 3339.

**Figure 12.**
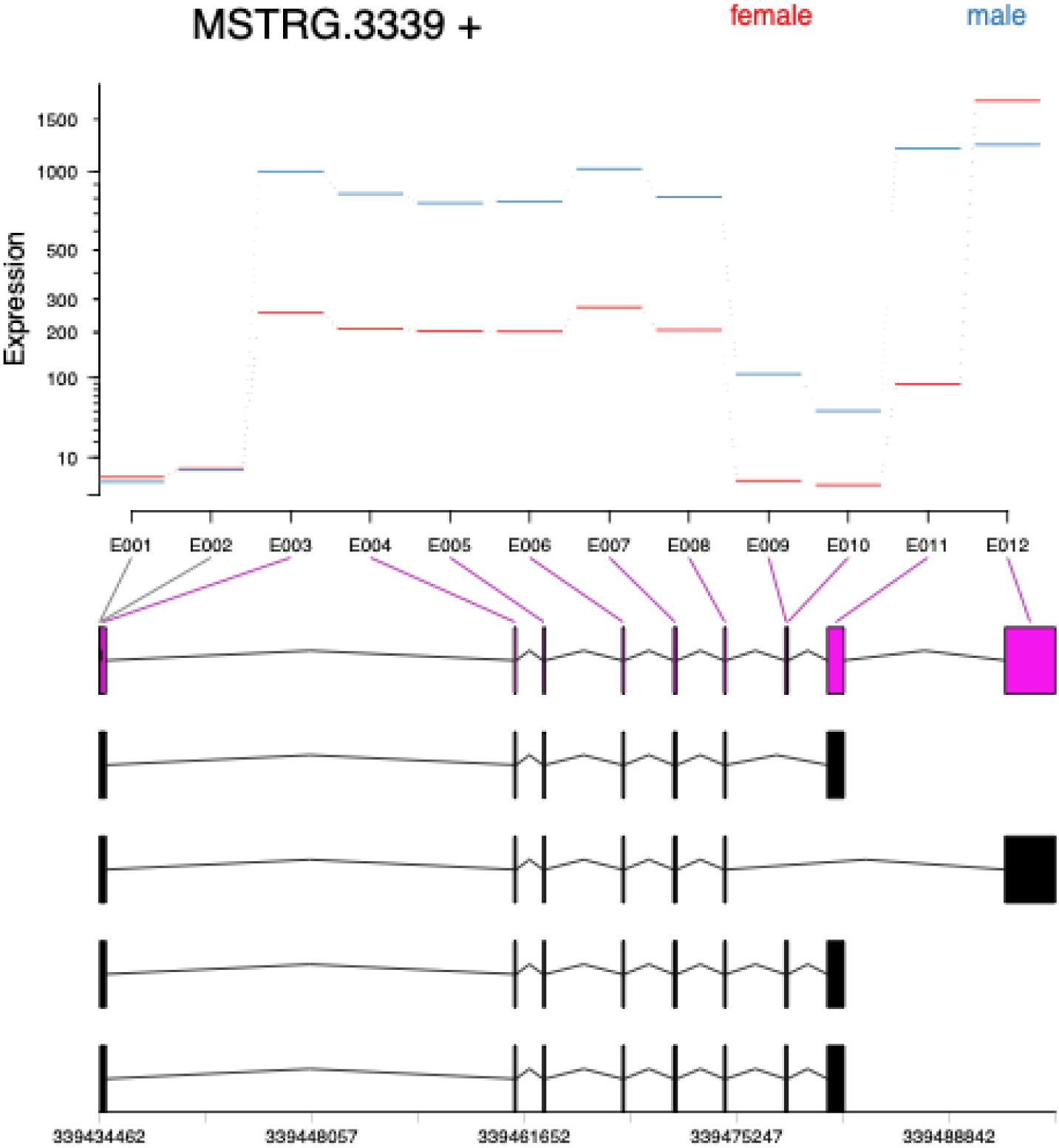
Exon plots analysis for MSTRG 3339 to elucidate sex specificity in *D. albipictus.* Exons were quantified, followed by differential usage analysis for the three ufMale compared to the three ufFem samples. Exons E009 and E010 show downregulation in male samples.

**Figure 13.**
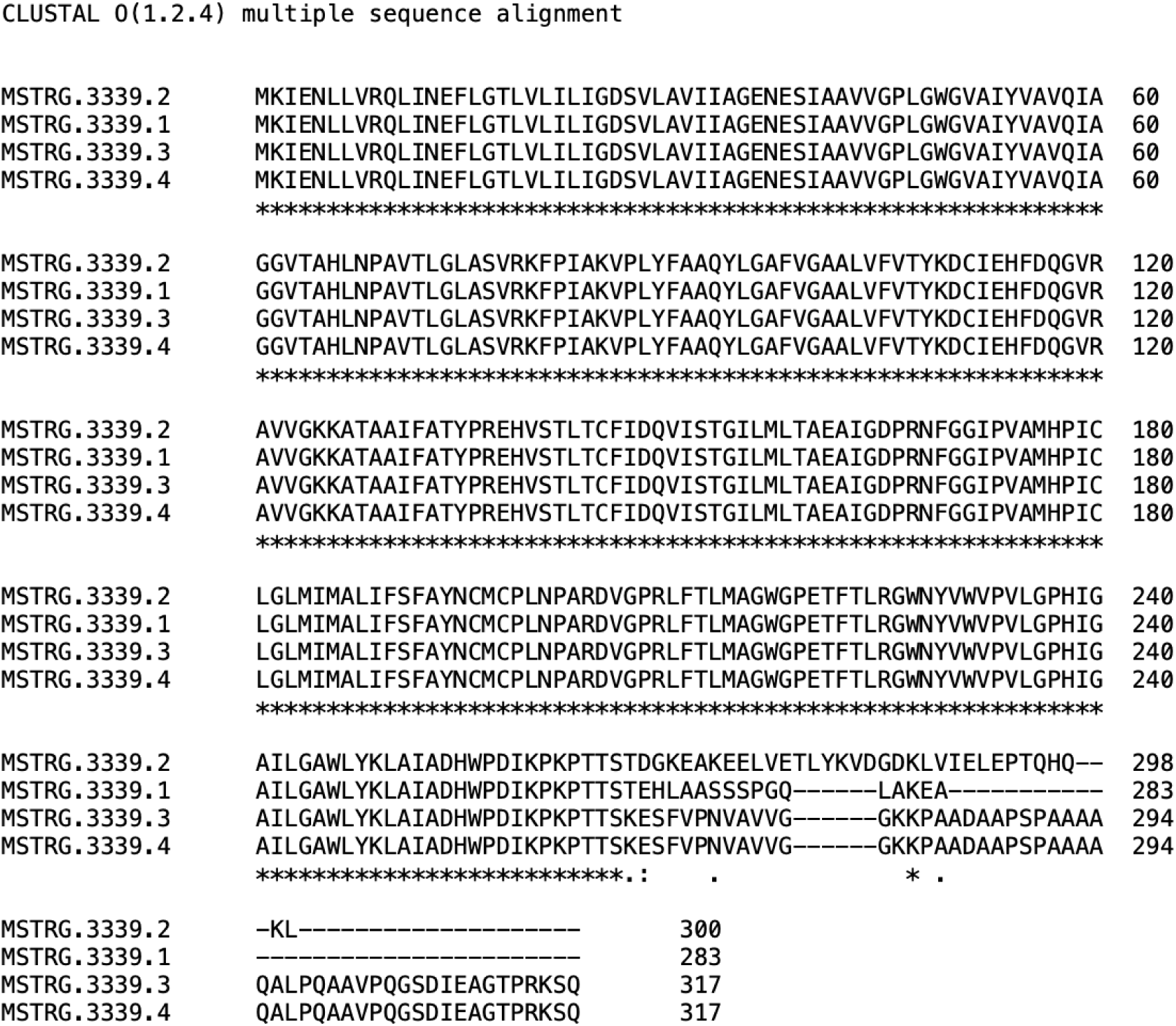
Protein alignment of MSTRG 3339 transcripts MSTRG.3339.1-3 using Clustal. 125 amino acid sequence is absent in the male transcripts MSTRG.3339.1-2.

**Table 28.**
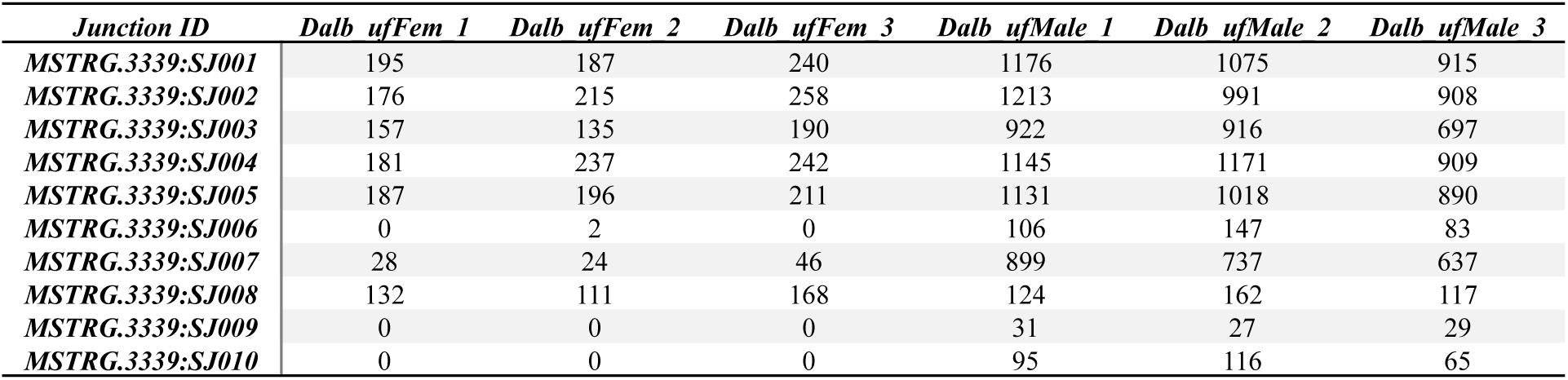
Splice junction counts for *D. albipictus* MSTRG 3339 showing sex-specificity of SJ006, SJ009, and SJ010 (male).

| *Junction ID* | *Dalb ufFem 1* | *Dalb ufFem 2* | *Dalb ufFem 3* | *Dalb ufMale 1* | *Dalb ufMale 2* | *Dalb ufMale 3* |
| --- | --- | --- | --- | --- | --- | --- |
| *MSTRG.3339: SJ001* | 195 | 187 | 240 | 1176 | 1075 | 915 |
| *MSTRG.3339: SJ002* | 176 | 215 | 258 | 1213 | 991 | 908 |
| *MSTRG.3339: SJ003* | 157 | 135 | 190 | 922 | 916 | 697 |
| *MSTRG.3339: SJ004* | 181 | 237 | 242 | 1145 | 1171 | 909 |
| *MSTRG.3339: SJ005* | 187 | 196 | 211 | 1131 | 1018 | 890 |
| *MSTRG.3339: SJ006* | 0 | 2 | 0 | 106 | 147 | 83 |
| *MSTRG.3339: SJ007* | 28 | 24 | 46 | 899 | 737 | 637 |
| *MSTRG.3339: SJ008* | 132 | 111 | 168 | 124 | 162 | 117 |
| *MSTRG.3339: SJ009* | 0 | 0 | 0 | 31 | 27 | 29 |
| *MSTRG.3339: SJ010* | 0 | 0 | 0 | 95 | 116 | 65 |

## Discussion

In this study, we generated high-quality transcriptome data for five *Dermacentor albipictus* sample types and used them to annotate the newly released *D. albipictus* genome. More than 79.95 million reads were collected per sample, with over 90% of reads successfully mapped and more than 85% uniquely mapped. This effort resulted in the identification of 47,811 transcripts and 22,680 genes, of which 47.88% (10,860) were annotated with GO functions and 58.39% (13,243) with PFAM IDs. Homologous genes in two closely related tick species, *D. andersoni* and *D. silvarum*, were also identified, providing a comparative framework for understanding gene function in *D. albipictus*. These results represent a significant step forward in characterizing the molecular biology of this tick species and lay the groundwork for future functional genomic studies.

Differential expression analyses identified overexpressed genes for five pairwise comparisons and GO enrichment analyses were used to highlight strongly affected biological processes. In embryos vs adults, embryos showed upregulation of genes involved in DNA repair, replication, and transcription regulation, reflecting their high mitotic activity and developmental needs (Adryan and Teichmann 2010). In contrast, adults exhibited enrichment for proteolysis, carbohydrate metabolism, and host-defense perturbation processes, which are likely linked to their feeding behavior and interaction with hosts (Perner et al. 2016; Gulia-Nuss et al. 2016). In males vs females, male samples displayed higher expression of peptidases and carbohydrate metabolism-related genes, while female samples had fewer differentially expressed genes but included some involved in translation (Obregón et al. 2019).

The male reproductive system compared to males showed enrichment for processes such as proteolysis, chromosome condensation, carbohydrate metabolism, and double-strand break repair, reflecting its role in spermatogenesis (Kiyozumi and Ikawa 2022; Sonenshine et al. 2011a). Peptidases (e.g., MSTRG.10729, MSTRG.10218) were strongly upregulated, alongside histamine-binding proteins (e.g., MSTRG.4914), which suppress host immune responses to enable prolonged blood feeding (Kazimírová and Štibrániová 2013). Additional enriched terms included steroid biosynthesis and ion transport, underscoring the metabolic demands of male reproduction.

In male reproductive system vs ovaries, male reproductive samples were enriched for proteolysis and carbohydrate metabolism, consistent with their role in spermatogenesis and male fertility. Ovaries showed upregulation of genes involved in DNA integration, DNA repair, and intracellular transport, reflecting their role in oogenesis and early embryonic development (Wang et al. 2021).

Comparing ovaries vs females, ovary-enriched genes were associated with DNA replication, intracellular transport, vesicle-mediated transport, and oogenesis-specific processes. In contrast, whole-body females showed high expression of histamine-binding proteins (e.g., MSTRG.19815, MSTRG.4914) and evasins that modulate host immune responses and maintain blood flow during feeding (Kazimírová and Štibrániová 2013). Blood coagulation inhibitors (e.g., MSTRG.6585, MSTRG.6479) were also overexpressed in females, highlighting their role in facilitating successful blood feeding (Kazimírová and Štibrániová 2013).

Another focus of this study was identifying candidate genes involved in sex determination pathways as little is known about these in the Acari. No homologs for genes involved in *Drosophila* sex-determination pathways have been identified in tick species, however homologs (*dsx1*,*dsx2*, *BTB1*, *BTB2*, *vg1, vg2, tra2 and ix*) have been identified in predatory mites, *Metaseiulus occidentalis* and *Neoseiulus cucumeris* (Zhang et al. 2019; Pomerantz and Hoy 2015). Not all of these genes undergo sex-specific splicing; however, *dsx1*, *dsx2,* and *BTB2* have male-biased expression.

Our global exon utilization analysis revealed widespread sex-specific alternative splicing events in *D. albipictus*, affecting 6,852 genes and over 26,000 exons - notably MSTRG.4611, a gene encoding a fatty acid CoA ligase-like protein with male-specific isoforms that may modify substrate specificity during spermatogenesis. Another example is MSTRG.10562, which encodes a ralA-binding protein 1-like protein with male-specific isoforms lacking a 125-amino-acid region, potentially influencing GTPase-mediated signal transduction. Additionally, MSTRG.3339 encodes an aquaporin-9 homolog with male-specific isoforms that alter the carboxyl terminus outside the membrane channel domain.

We identified four putative doublesex (*dsx*) homologs in *D. albipictus*, two of which showed strong sex-specific expression or splicing patterns, MSTRG.10335 and MSTRG.10337. MSTRG.10335 is expressed in both sexes but displays sex-specific splicing patterns, with distinct isoforms present in males and females. Notably, this represents the first documented case of sex-specific splicing of a doublesex-like gene in chelicerates, a mechanism previously thought to be absent in this group (Price, Egizi, and Fonseca 2015). Such splicing patterns are reminiscent of *dsx* regulation in insects like *Drosophila melanogaster (Burtis and Baker 1989)*, *Aedes aegypti (Salvemini et al. 2011)*, and *Anopheles gambiae (Scali et al. 2005)* where alternative splicing is central to sexual differentiation. The discovery of sex-specific splicing in ticks suggests this regulatory mechanisms may have existed within arthropods but was lost or modified in other chelicerate lineages (Price, Egizi, and Fonseca 2015). Additionally, MSTRG.10337 exhibits male-specific expression, making it a promising candidate for further investigation as a potential sex determination regulator.

The identification of these candidates is significant because *dsx* has been central to developing genetic control tools such as pgSIT and gene drives in insects. For example, targeted knockouts of female-specific isoforms of *dsx* in *Anopheles gambiae* result in intersex individuals incapable of blood feeding or reproducing (Kyrou et al., 2018). Similarly, pgSIT systems leveraging sex-specific splicing have been successfully developed for *Drosophila melanogaster* (Kandul et al. 2019) and *Aedes aegypti* (Li et al., 2024) and have been successfully utilized in genetic sex-sorting systems (Weng et al. 2023) (Weng et al. 2024). These approaches could potentially be adapted for ticks by targeting sex-specific splicing events in MSTRG.10335. Functional validation through CRISPR or RNAi experiments will be necessary to confirm the roles of these genes in sex determination.

For vector control purposes, identifying transcriptional regulatory elements expressed within the germline is an important step. RNA sequencing data from male and female reproductive systems will provide insights into sex specific gene regulation which has been demonstrated in other vector species (Akbari et al. 2014). Additionally, for control approaches relating to fertility, it is important to identify gene targets linked to sex determination, spermatogenesis, and oogenesis.

In *D.variabilis,* Neprilysin (membrane metalloprotease), and ferritins (iron storage) have been identified as highly expressed in male reproductive organs (Sonenshine et al. 2011b). Neprilysin, which plays an important role in the function of sperm and fertilization in *D.melanogaster* (Sitnik et al. 2014), may be an interesting gene to study further, as would Ferritins, which are also important in reproduction, evidenced by knockdown studies in *Ixodes ricinus* showing reduced oviposition in females (Hajdusek et al. 2009). Alternative potential targets include the male-produced peptides Efα and Efβ and vitellogenin receptors (VgRs). Efα and Efβ have been shown to trigger female *Amblyomma hebraeum* to fully engorge post male insemination (Weiss and Kaufman 2004). VgRs are key to the production of functional yolk proteins in oocytes, which allow the development of embryos. They have been identified as potential targets for vaccine development (Roe et al. 2008; Meng and Sluder 2018), in addition to potential uses in pathogen transovarial-transmission blocking. *Babesia spp* (*B. gibsoni* and *B. bovis*) have been demonstrated to be blocked from transmission to oocytes via the silencing of VgR in *Haemaphysalis longicornis* (Boldbaatar et al. 2008) and *Rhipicephalus microplus* (Hussein et al. 2019), as well as interfering with oocyte development and ovary maturation (Smith and Reuben Kaufman 2013). In *Amblyomma hebraeum*, knockdown of VgRs results in delayed development of ovaries and oviposition. Furthermore, VgRs have also been identified in *D. variabilis* and *R. microplus*, and are transcribed differentially across life stages, sex, and tissues (R. D. Mitchell 3rd et al. 2007; Seixas et al. 2018). New genome and transcriptome data will aid in the identification of VgR homologs within this *D. albipictus*.

The availability of new *D. albipictus* transcriptome data presents an advancement in understanding the molecular mechanisms underlying key biological processes in this tick species. By analyzing the transcriptome, researchers can identify and characterize genes that play crucial roles in various physiological pathways, including those related to reproduction and acaricide resistance. Overall, the integration of transcriptome data with genetic and functional studies holds promise for advancing the development of effective control strategies for *D. albipictus* and mitigating its impact on wildlife, livestock, and public health.

## Methods

### Tick Rearing

The *D. albipictus* colony (Rhodes 1998 strain) used in this study was propagated on cattle at the Knipling-Bushland US Livestock Insects Research Laboratory (KBUSLIRL; Kerrville, Texas USA). Larval and engorged female *D. albipictus* are incubated in glass aquaria with temperature and humidity maintained at 24℃, 92-97% RH. Humidity was regulated with a potassium nitrate supersaturated salt solution at the base of the aquarium. A bovine calf was infested with *D. albipictus* larvae under a muslin cloth patch, which was adhered to the shaved region of the animal using a veterinary-approved adhesive. Females that fed to repletion and dropped from the host were collected, and ovaries from pre-oviposition females were dissected 10 days post-drop. For collection of egg masses, engorged females (n = 20) were placed dorsal side down on double-sided sticky tape adhered to the interior of a Petri dish to await oviposition. At the onset of oviposition, eggs deposited over a 6-hour period were collected, snap frozen in liquid nitrogen, macerated in Monarch DNA/RNA Protection Reagent (NEB), and stored at -80℃. Unfed adult females and males were obtained from the host at 18-19 days after larval infestation, which coincides with the nymphal molt to adults. For the collection of reproductive systems from fed, unmated males, newly molted adults were obtained from the host and transferred to a separate, male-only patch on the same host. The males attached and were allowed to feed for 4 days, after which they were pulled from the host for dissection. All dissected tissues were transferred to Monarch DNA/RNA Protection Reagent, macerated, and stored at -80℃ before extraction. Three biological replicates of each sample type were analyzed, yielding a total of 15 samples.

### RNA Extraction

Illumina RNA sequencing was conducted to quantify gene expression patterns. RNA was extracted from all life stages and tissues using the Monarch Total RNA Miniprep Kit (NEB), following the manufacturer’s protocol.

After RNA extraction, DNase treatment was conducted using RNase-free DNase I. The integrity of the RNA was assessed, and mRNA was isolated from the total RNA. RNA-seq libraries were prepared for Illumina sequencing using the NEB library prep kit. Library sequencing was conducted using Illumina NextSeq 2000 to give paired-end reads of 150nt and 20 million reads per library. RTA1.18.64 was used for base calls and converted to FASTQ using bclfastq 1.8.4.

### Transcriptome Assembly and Annotation

RNA sequencing reads were trimmed to remove adapter sequences with AdapterRemoval (Schubert, Lindgreen, and Orlando 2016) and aligned to the GCA_038994185.2 genome with STAR (Dobin et al. 2013) using the 2-pass method. The 2-pass method enables more accurate quantification of spliced reads in the absence of annotations. The aligned reads for each sample were processed separately with StringTie (Pertea et al. 2015) to generate de novo transcript annotations, which were subsequently merged, with StringTie, to generate the final GTF file Dalb_finalv2.merged.trimmed_reads.gtf.

Transcript coordinates defined in Dalb_finalv2.merged.trimmed_reads.gtf were used to generate transcript sequences. Encoded ORFs were predicted using ORFfinder (Sayers et al. 2012) and the longest one for each transcript was parsed. Protein domains were identified with HMMSCAN (Eddy 2011) and GO terms associated with PFAM domains were added using pfam2GO (A. Mitchell et al. 2015) mappings. Protein and transcript homologs in two tick species were identified by blast searches against proteomes and transcriptomes of *Dermacentor andersoni* (GCF_023375885.1) and *Dermacentor silvarum* (GCF_013339745.2).

### Gene Quantification, Differential Expression and GO Enrichment Analyses

Genes defined in Dalb_finalv2.merged.trimmed_reads.gtf were quantified with featureCounts (Liao, Smyth, and Shi 2014) and the count values were then transformed into TPM (Transcripts Per Million) and FPKM (Fragments Per Kilobase of transcript per Million mapped reads) values with a Perl script. TPM values were used to perform PCA and clustering analyses. Plots were generated in R using ggdendro and ggplot2 packages (Kassambara 2015).

Differential analysis runs for five pairwise comparisons (embryo vs adult; male vs female; mrs vs male; ovary vs female, mrs vs ovary) were conducted with DESeq2 (Love, Huber, and Anders 2014). The identified differentially expressed genes were used to perform GO enrichment analyses for the biological process (BP) ontology terms using the topGO (Rahnenfuhrer 2024) R package. Up- and down-regulated gene sets were analyzed separately using the weight01 algorithm and Fisher’s exact test.

### Sex-specific Splicing and Doublesex Candidate Identification

Differential exon usage analyses were performed with DEXSeq (Anders et al. 2012). Briefly, a flattened GFF file was generated with dexseq_prepare_annotation.py and exon parts were quantified with dexseq_count.py scripts provided by the DEXSeq package. Exon plots were generated with the plotDEXSeq function. *Doublesex* candidates were identified by parsing blast results that showed homology to genes annotated as *dsx* or *dsx-like* in *D. andersoni* and *D. silvarum*. Splice junction counts were extracted from SJ.out.tab files generated by the STAR aligner. Transcript structures and splicing patterns were visualized in Integrative Genomics Viewer (IGV) (Robinson et al. 2011).

### Data Availability

The Illumina sequencing reads for the *D. albipictus* embryos, male reproductive system, ovaries preoviposition, unfed males and unfed females have been submitted to NCBI Sequence Read Archive within the Transcriptome database (BioProject Accession Primary PRJNA1072088; RNA-Seq SRA study SRP492691, SRR28628997, SRR28628998, SRR28628999, SRR28629000 and SRR28629001).

### Ethical Conduct of Research

All animals were handled in accordance with the Guide for the Care and Use of Laboratory Animals as recommended by the National Institutes of Health and approved by the UCSD Biological Use Authorization (BUA #R2401), and animal procedures were approved by the Institutional Animal Care and Use Committee of the KBUSLIRL (IACUC Protocol # 2023-13).

## Acknowledgments

We thank Wayne Ryan (KBUSLIRL) for helping with tick husbandry and Greta Buckmeier and Deanna Bodine (KBUSLIRL) for laboratory assistance. This work was supported by funding from the Bill & Melinda Gates Foundation (INV-056448) awarded to O.S.A. The views, opinions, and/or findings expressed are those of the authors and should not be interpreted as representing the official views or policies of the U.S. government.

## Author Contributions

O.S.A., and I.A. conceptualized the study. P.U.O., and P.S. performed all sample preparations and sequencing; I.A. performed transcriptome annotation; I.A., and R.T.M.E, performed data analysis; R.T.M.E., and E.H wrote the first draft of the manuscript. All authors contributed to analyzing and compiling the data. All authors contributed to writing and approving the final manuscript.

